# Psychiatric Genomics: An Update and an Agenda

**DOI:** 10.1101/115600

**Authors:** Patrick F Sullivan, Arpana Agrawal, Cynthia M Bulik, Ole A Andreassen, Anders D Børglum, Gerome Breen, Sven Cichon, Howard J Edenberg, Stephen V Faraone, Joel Gelernter, Carol A Mathews, Caroline M Nievergelt, Jordan Smoller, Michael C O’Donovan, for the Psychiatric Genomics Consortium

**Author notes:** Correspondence to: Dr Sullivan. Submitted to: *American Journal of Psychiatry (Review).

## Abstract

The Psychiatric Genomics Consortium (PGC) is the largest consortium in the history of psychiatry. In the past decade, this global effort has delivered a rapidly increasing flow of new knowledge about the fundamental basis of common psychiatric disorders, particularly given its dedication to rapid progress and open science. The PGC has recently commenced a program of research designed to deliver “actionable” findings - genomic results that (a) reveal the fundamental biology, (b) inform clinical practice, and (c) deliver new therapeutic targets. This is the central idea of the PGC: to convert the family history risk factor into biologically, clinically, and therapeutically meaningful insights. The emerging findings suggest that we are entering into a phase of accelerated translation of genetic discoveries to impact psychiatric practice within a precision medicine framework.

**Collaborators:** PGC Coordinating Committee: Mark Daly, Michael Gill, John Kelsoe, Karestan Koenen, Douglas Levinson, Cathryn Lewis, Ben Neale, Danielle Posthuma, Jonathan Sebat, and Pamela Sklar.

## Introduction

Heredity is intimately related to the history of psychiatry. Clinical observations by early physicians noted the tendency of mental illnesses to run in families. In the 20^th^ century, these anecdotes were systematically evaluated and some were confirmed in rigorous twin, family, and adoption genetic epidemiological studies. This exceptional body of evidence provided a major etiological clue for the field: common psychiatric disorders have a moderate to strong tendency to run in families in large part due to genetic inheritance (1, 2).

We now know that these genetic effects are relatively subtle and non-deterministic: most people with a strong family history are not themselves affected. Moreover, most psychiatric disorders do not “breed true” with increased risks spread across multiple disorders. In fact, the diverse clinical manifestations and variable course observed for many common psychiatric disorders are consistent with complex and relatively subtle genetic effects. For adult-onset common psychiatric disorders in particular, development is often within normal limits, and many experience relapsing/remitting illnesses with preservation of basic neurological function but often with some degree of erosion of higher components of cognition.

In the last decade, it has become technically and economically feasible to interrogate the genome directly with increasing resolution and completeness. Instead of studying the heredity of psychiatric disorders in a general way (e.g., via studies of pedigrees, twins, or adoptees), we can now evaluate the genomes of cases and controls at several levels of precision quickly and inexpensively. Indeed, heritability itself can be assessed directly from genome-wide genetic data (3, 4).

By carefully evaluating the successes and failures of psychiatric genetics in the past three decades, we now have a solid fix on how to dissect the “family history risk factor” into far more precise and mechanistic components. We can identify genetic variants that contribute to risk, and are moving toward understanding the mechanisms by which they act. The field has learned an enormous amount, and is poised to make fundamental advances that could profoundly improve understanding.

In 2009, the Psychiatric Genomics Consortium (PGC) published three foundational papers regarding genome-wide association studies (GWAS) (5-7). GWAS is a genomic study design that focuses on the impact of common genetic variation in almost all genes in the human genome. The initial PGC papers covered the core concepts, history, the rationale, genomic assays, statistical analysis, interpretative framework, and the importance of cross-disorder studies in psychiatry. Full background of the terminology, core concepts, and strategy of GWAS can also be found in these papers. Basic terms are defined in *Table S1.*

This review provides an update on what we have learned, and puts forth an agenda for the next five years. A key lesson was the need for a global community effort in psychiatric genetics because the required samples sizes are far beyond the reach of any single group. To enable these studies, in 2007 we formed the PGC (URLs). Our overarching goal is to deliver actionable knowledge, i.e., genetic findings whose biological implications can be used to improve diagnosis, develop rational therapeutics, and craft mechanistic approaches to primary prevention.

## Clarity in retrospect

A key unknown was genetic architecture, particularly the sizes of the underlying genetic effects. A decade ago, these were unknown and subject to considerable speculation with hypotheses suggesting that genetic discovery for psychiatric disorders would be anywhere from highly tractable to impossible. If the genetic effects were few, common, and large, relative modest sample sizes would be sufficient. A few early studies hinted that relatively small samples might suffice (e.g., the large effects of *APOE* on Alzheimer’s disease or *CFH* on age-related macular degeneration) (8, 9), and these may have led to expectations that gene discovery would be straight-forward.

The power calculations are not difficult: for a given number of cases and controls (plus assumptions of allele frequency, genetic model, significance threshold, and power), it is easy to compute the minimum detectable genotypic relative risk (GRR). For example, *Figure 1a* shows the 90% power curve for a GWAS of 1,000 cases and 1,000 controls.

**Figure 1a.**
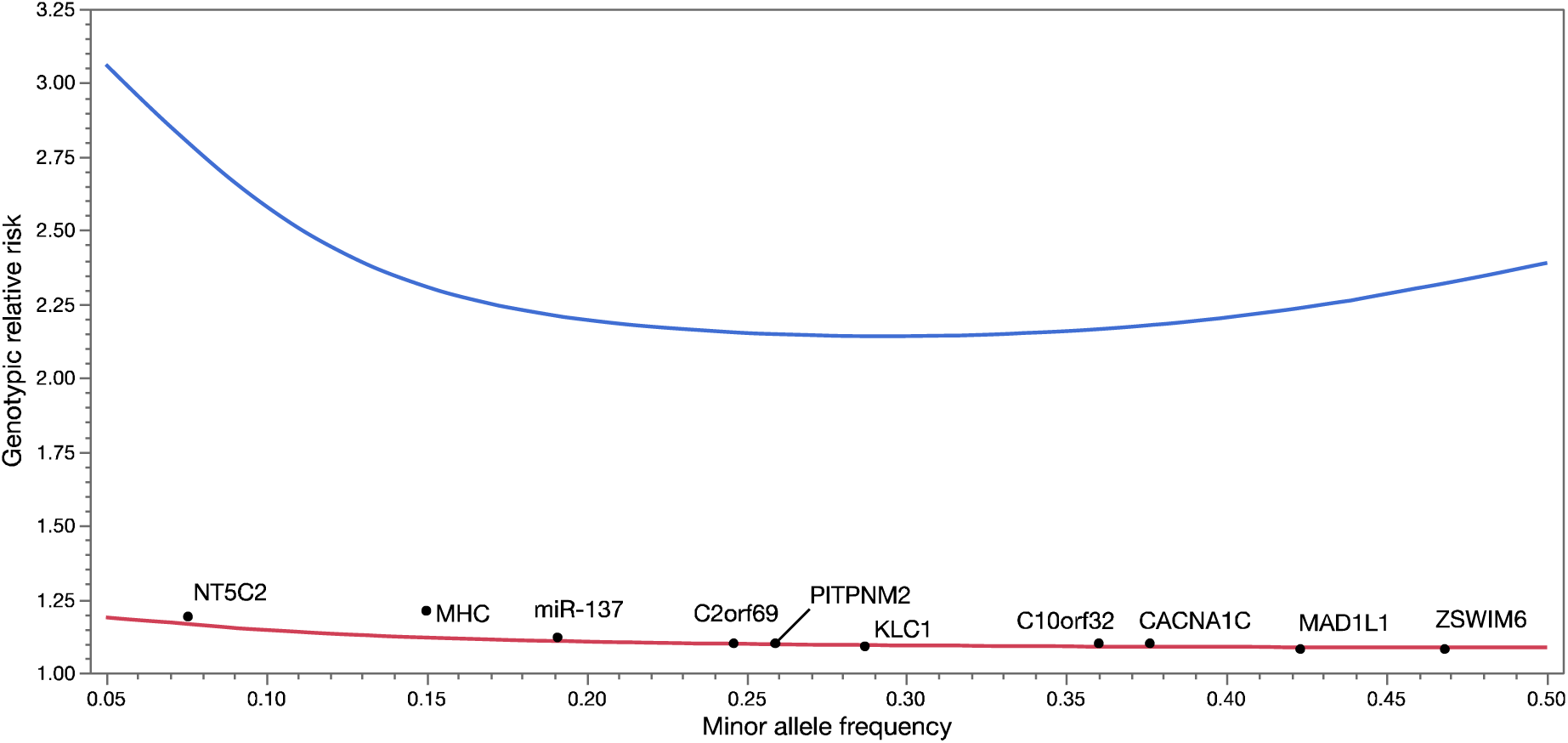
Statistical power. Upper curve (blue) shows minimum detectable genotypic relative risks for common variants for 1,000 cases and 1,000 controls (90% power, additive model, lifetime risk 0.01, α=5e-8). Lower curve (red) shows 90% power for the PGC 2014 schizophrenia paper (37,000 cases and 113,000 controls, additive model, lifetime morbid risk 0.01, α =5e-8). Black dots show the top 10 loci in the PGC schizophrenia report. These loci are highly significant with P-values ranging from 1.7e-13 to 3.8e-32.

Like most investigators in human complex disease genomics, we had limited data to allow us to narrow bounds on the search space. We quickly learned that optimistic assumptions of large genetic effect sizes for these disorders were incorrect. The initial GWAS for psychiatric disorders had sample sizes ^~^1,000 cases enabling excellent power to detect GRR ≥2.5. However, these effects were not found for schizophrenia (10), bipolar disorder (11), major depressive disorder (12), or ADHD (13). *Figure 1a* also shows the 90% power curve for the largest published study of any psychiatric disorder (37,000 cases) (14), and only two of 128 independent loci had GRR ≥1.2. Compellingly, we can now demonstrate that common genetic variants with GRR above ^~^1.24 for schizophrenia can be excluded with ^~^100% power.

Genetic effects that are common and large are unusual for human diseases and traits studied using GWAS (*Figure 1b*). They are occasionally found for less complex and well-measured conditions (e.g., infectious diseases, rare adverse drug reactions, and eye diseases). To our knowledge, the largest common genetic variant associations observed to date in psychiatry are for alcoholism in people of East Asian ancestry (GRR ^~^6.2) and clozapine-induced agranulocytosis (GRR ^~^5.3) (15, 16).

**Figure 1b.**
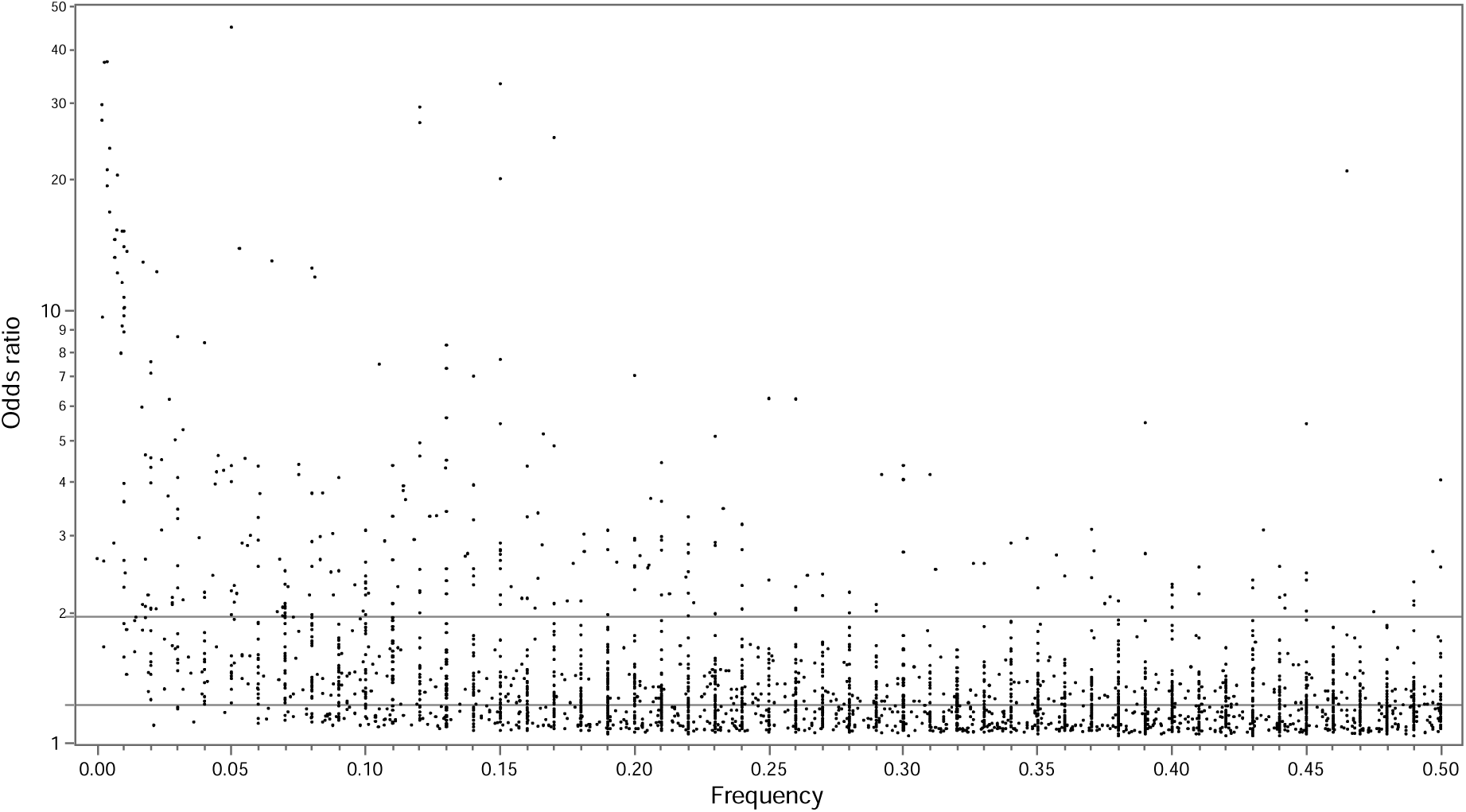
Odds ratios (OR, log_10_) and allele freguencies from published GWAS. From EBI-NHGRI GWAS catalog (URLs, accessed 1/27/2017), contains 2,308 GWAS published 3/2005-7/2016. There are 9,485 SNP-trait associations (P ≤ 1e-8) including 7,487 SNPS and 870 traits. Dots show freguency and OR (OR transformed to be >1 and allele freguencies to 0-0.50). Contours show densest areas of the plot. Horizontal lines show 50^th^ (1.22) and 90^th^ (1.95) percentiles for ORs: most associations are subtle. Of 62 associations with 0R>5, most are for infectious disease (31; e.g., influenza susceptibility), pharmacogenomic (13; e.g., rare adverse drug reactions like flucloxacillin-induced liver injury), eye disease (4; e.g., glaucoma), or pigmentation (2; e.g., blue vs. brown eyes). Only a few diseases have atypically large ORs (e.g., celiac disease, melanoma, membranous nephropathy, myasthenia gravis, ovarian cancer, Parkinson’s disease, progressive supranuclear palsy, thyrotoxic hypokalemic periodic paralysis, and type 1 diabetes). The only psychiatric finding was alcohol consumption and ALDH2 in individuals of East Asian ancestry.

## Genetic architecture and models of disease

Elucidation of the genetic architecture underlying these disorders is the major goal of the PGC. How many susceptibility or protective variants are there? What are their frequencies and effect sizes? How do they exert their effects? Do these variants interact with one another or with environmental risk factors? Crucially for biological understanding, which genes are affected by these variants?

It is heuristically useful to consider the bookends. The extreme models are that psychiatric disorders are caused by (a) the cumulative impact of hundreds or thousands of common genetic variants each of subtle effect (common-disease/common-variant model) or (b) many different gene-disrupting variants of strong effect (multiple rare variant model). In the latter model, every person with a serious psychiatric disorder would have a strong effect variant and these would cluster in a set of genes important to brain development and function.

These models were passionately debated. Some authors expressed profound hope that the multiple rare variant model was broadly explanatory (17-19). Others favored a common disease/common variant model, arguing that psychiatric phenotypes are comparatively subtle. Most investigators were agnostic. The PGC wished to design studies that would be informative whatever the underlying model (5).

## The initial strategy of the PGC

A consistent lesson from the history of psychiatric genomics was that these are very hard problems: any search is going to be far more difficult than anticipated. Although we were hopeful that the initial GWAS might deliver insights, we created the PGC in order to hedge our bets: we needed a framework to aggregate data across studies with exceptional care and rigor if we were to progress. A critical step was to convince all groups that sharing *individual* data was essential – this is a foundational principle of the PGC and allows optimal quality control and analysis.

Moreover, to ensure progress, an “open-science” perspective was required. Genome-wide summary statistics of all PGC analyses are available for widespread use (URLs), and the vast majority of PGC genotype data that can be deposited are available to qualified researchers in a controlled-access repository. We recently have made available a list of the top loci from PGC analyses (both published and in preparation).

These early strategic decisions proved important: results from the first wave of psychiatric GWAS, circa 2008, were unimpressive. Although we were careful not to hype GWAS (5, 6), some prominent commentators voiced strong doubts about its value – even though careful review of the early results showed unequivocal indications of genetic effects. The first wave studies were simply underpowered, and combining studies to increase power was logical. Nevertheless, we persisted, and a 2012 letter signed by 96 psychiatric genetics investigators (“Don’t give up on GWAS”) anticipated the utility of GWAS should sample sizes increase (20).

To date, the PGC has published 24 main papers and 51 secondary analysis papers (*Table S2*). At least 141 additional papers have made use of PGC results. Many PGC papers are highly cited, but chief among them is the schizophrenia *Nature* (14) report which ranks among the most highly cited papers in 2014. The PGC is among the leading genomic consortia worldwide for open science and data sharing. These successes are a testimony to the fact-based strategy and persistence of the PGC.

## An update

What have we learned? We now have a sizable body of empirical results relevant to the common “versus” rare variant debate. All common psychiatric disorders with sufficiently large samples have a predominant common-disease/common variant contribution (21-23). Indeed, this is widely seen across human complex diseases like type 2 diabetes mellitus (24), and anthropometric traits like height (25) and body mass (26). Demonstrating a major role of common genetic variation on risk for human complex traits (including psychiatric disorders) is so widely and consistently documented that it is no longer particularly newsworthy.

There is a variable contribution of rare variation of strong effect. This tends to be larger for early onset, severe disorders and lesser for disorders with normal-range developmental trajectories and adult onset (*Figure 2a*). However, even for psychiatric disorders with many proven examples of rare variants of strong effect (e.g., intellectual disability or early-onset Alzheimer’s disease), there is always a contribution of common variation. Rare variant studies have proven more difficult than anticipated: to confidently identify rare variants of strong effect in typical clinical samples requires very large sample sizes, perhaps as many as ^~^100K cases (27). The protein-coding parts of the genome are replete with inconsequential variation, and current ways to predict functional consequences are imprecise (28). There is a lot of noise, and the signal is sparser and weaker than anticipated.

**Figure 2a.**
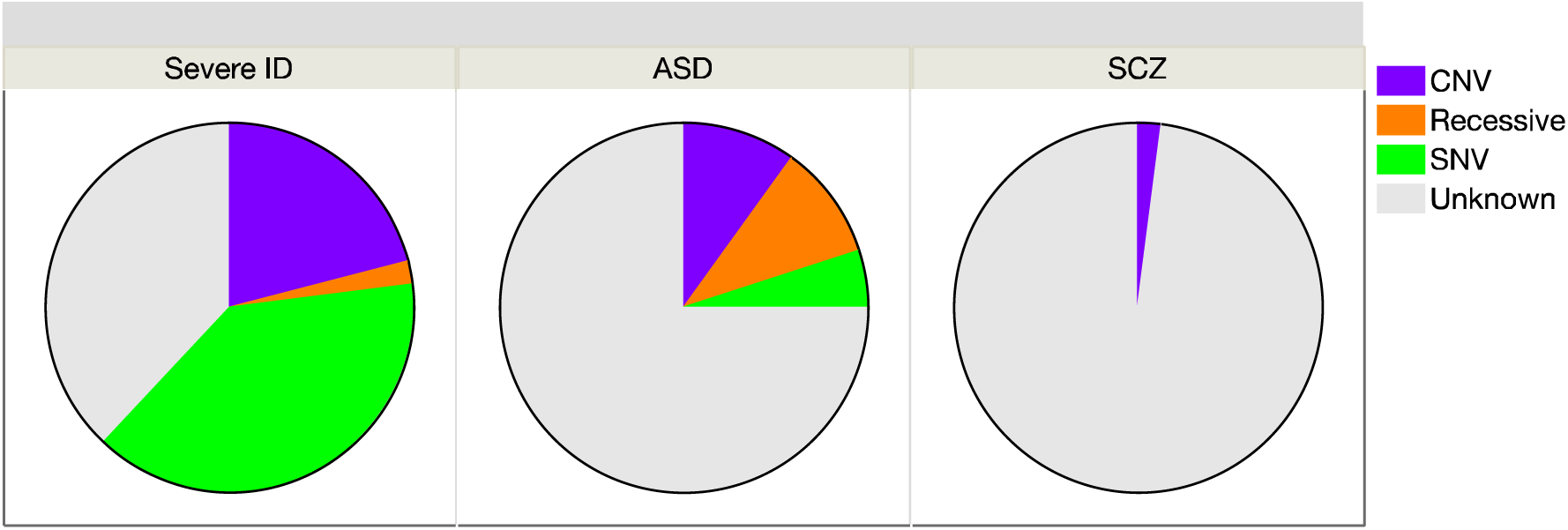
Genetic causes of severe intellectual disability (ID) (50), autism spectrum disorder (51, 52), and schizophrenia (53), including copy number variation (CNV), inherited known recessives, and single nucleotide variants (SNV). For severe ID, most SNV and CNV are de novo. The unknown grouping includes common variation, undiscovered rare genetic causes, phenocopies, and causation due to non-genetic effects.

*Table 1b* shows current sample sizes and notable findings for the nine PGC working groups. Schizophrenia has accumulated the most data for both common and rare variation. *Figure 2b* shows significant results from GWAS, copy number variation (CNV), and exome sequencing studies (14, 29, 30). Most findings are for common variation. Multiple rare CNVs have been implicated; most are multi-genic and all increase risk for several psychiatric disorders and neurological diseases (29). *SETD1A* is the only gene implicated to date by whole exome sequencing studies (30), but other such studies have only found hints of biological pathways by focusing on extremely rare variation (31, 32). It was widely anticipated that exon variation in the 0.005 to 0.01 allele frequency range would be readily found but this has not been observed (33), and a recent study of height required over 700,000 subjects to identify loci in this range (34). In a direct comparison, common variation had 14-28 times more impact on risk for schizophrenia than rare CNVs or rare exonic variation (35).

**Figure 2b.**
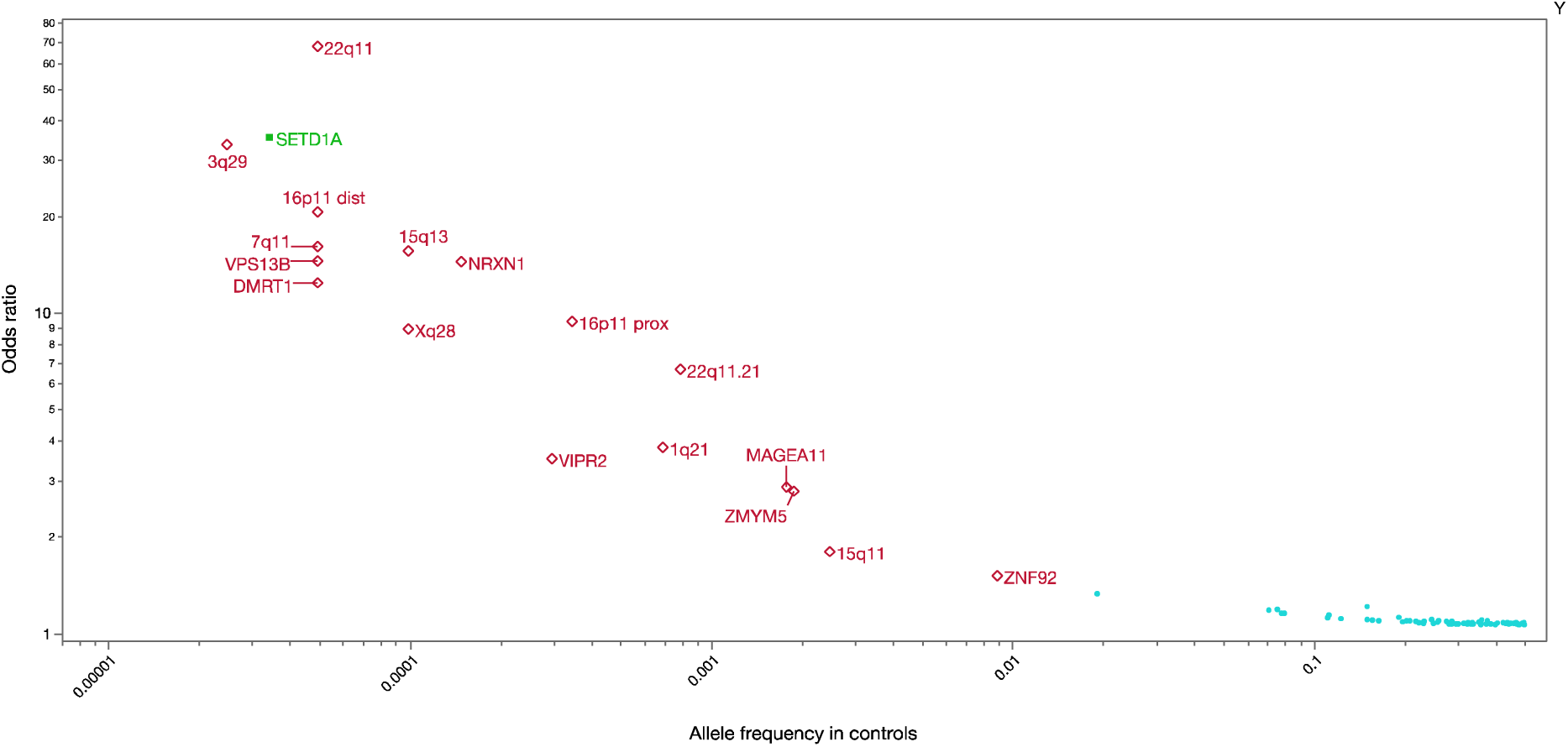
Significant genetic associations for schizophrenia. Y-axis is log_10_ of odds ratio. X-axis is log_10_ of allele freguency in controls. Odds ratios transformed to be >1 and freguencies to be ≤ 0.5. The dots on the lower right ( cyan ) shows common-variant associations for schizophrenia (P<1e-8) (14). Open diamonds (red) show copy number variation associated with schizophrenia (29). Filled sguare (green) shows the lone variant identified using whole exome seguencing (30).

Another major finding has been the repeated empirical documentation of important genetic overlap (particularly common variation) between most or all adult- and childhood-onset psychiatric disorders (21, 22). It is clear that psychiatric nosology has not “carved nature at the joints”. Moreover, the common variant genetic architecture of many disorders blends into normal phenomena. For example, there are sizable genetic correlations of MDD with personality traits like neuroticism and easily-assessed depressive symptom measures. Other findings suggest reconceptualizations may be needed. For example, anorexia nervosa had a significant positive genetic correlation with schizophrenia, significant negative genetic correlations with body mass index and unfavorable metabolic measures, and significant positive genetic correlations with favorable metabolic factors. This pattern of findings suggests that the roots of anorexia may be not only psychiatric, but also metabolic in origin.

This is an evolving area with regular increases in confident knowledge. To encourage rapid dissemination of results, the PGC regularly compiles an shares a list of the strongest findings for the disorders it studies (URLs).

## PGC, an agenda

Attempts to understand the genetic basis of psychiatric disorders – to untangle and concretize the family history risk factor – have never been easy. However, by incorporating empirical results, a data-driven and logical way forward has emerged, and it is likely that these efforts will continue to yield important new knowledge. Many groups are active in this area, but the PGC has emerged as the key umbrella organization for a large portion of this work. A basic description of the PGC and its core principles is given in *Table 1a.* Key technical aspects include its dedication to rigorous methodologies and its stance as a “mega-analysis” consortium with PGC members sharing individual-level genotype and phenotype data.

**Table 1a.**
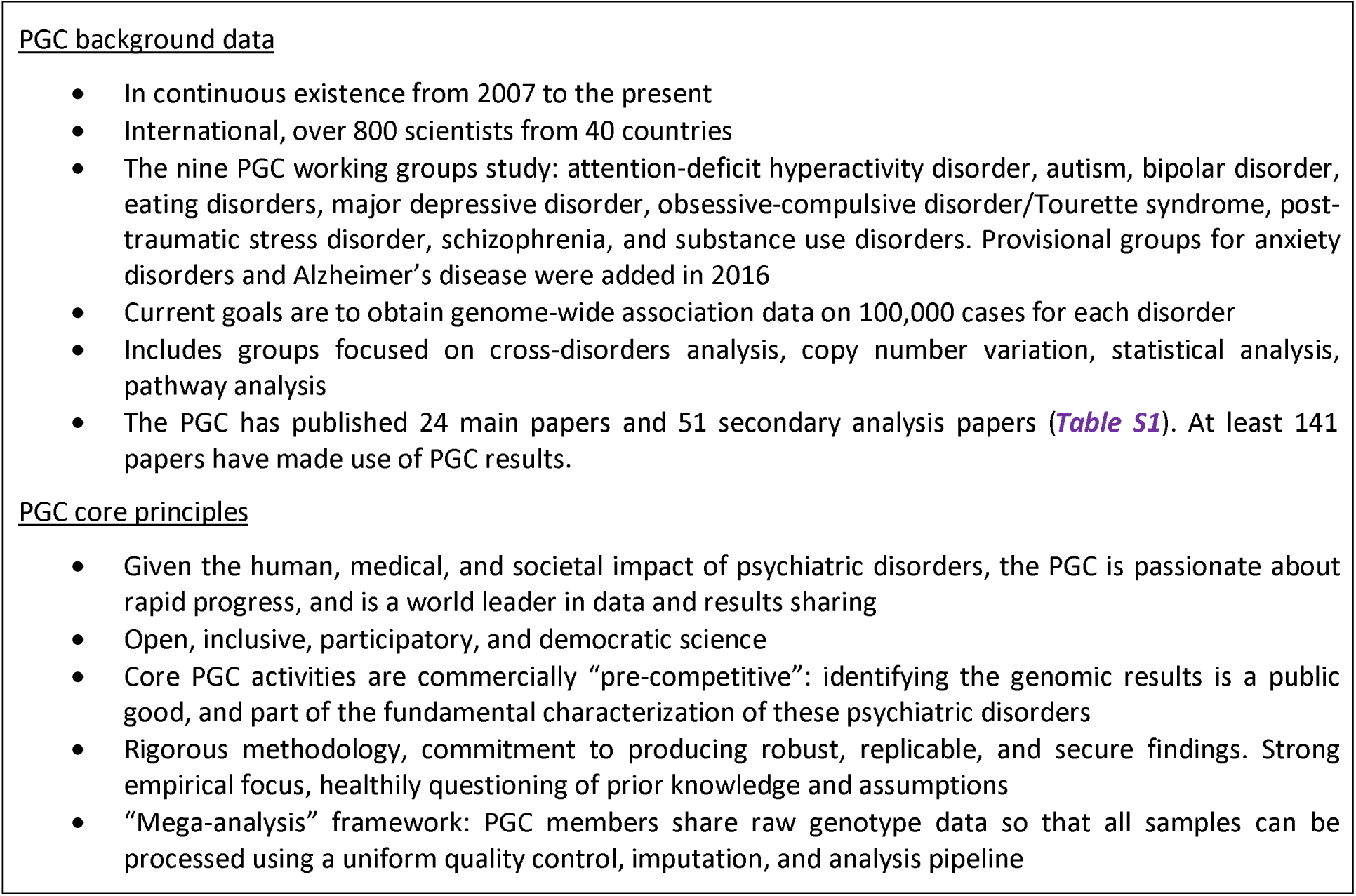
PGC background information and core principles (URLs).

With continued support from the NIMH (and new support from NIDA), the PGC recently initiated a program of research designed to deliver “actionable” findings, genomic results that (a) reveal the fundamental biology, (b) inform clinical practice, and (c) deliver new therapeutic targets. This is the central idea of the PGC: to convert the family history risk factor into biologically, clinically, and therapeutically meaningful insights. This program of research has six aims, three focused on common variation and three on rare variation (*Table 1c*).

**Table 1b.** Update, current findings for each PGC working group. Ncase=number of cases. Hits=independent associations reaching genome-wide significance. Twin-h^2^=heritability estimated from twin studies. SNP-h^2^=heritability estimated from GWAS results.

| PGC group | N<sub>case</sub> | Hits | Twin-h<sup>2</sup> | SNP-h<sup>2</sup> | Strongest genetic correlation |
| --- | --- | --- | --- | --- | --- |
| Schizophrenia | 60,995 | 155 | 81% | 45% | Bipolar disorder |
| MDD | 130,664 | 20+ | 30-40% | 14% | Neuroticism |
| Bipolar disorder | 20,352 | 19 | 80% | 21% |  |
| ADHD | 20,183 | 12 | 70-80% | 22% | Educational attainment |
| Autism spectrum disorder | 18,381 | 3 | 75% | 12% | Subjective well-being |
| Anorexia nervosa | 3,495 | 1 | 56% | ~20% | Metabolic traits |
| Substance use disorders | 3,772 | 1 | 50% | ~10% | Smoking |
| Tourette syndrome | 4,232 | 1 | 60-80% | 58% | OCD |
| OCD | 2,688 | 0 | 45-65% | 37% | Tourette syndrome |
| PTSD | 3,749 | 0 | 30-40% | 5-35% | Schizophrenia |

**Table 1c.**
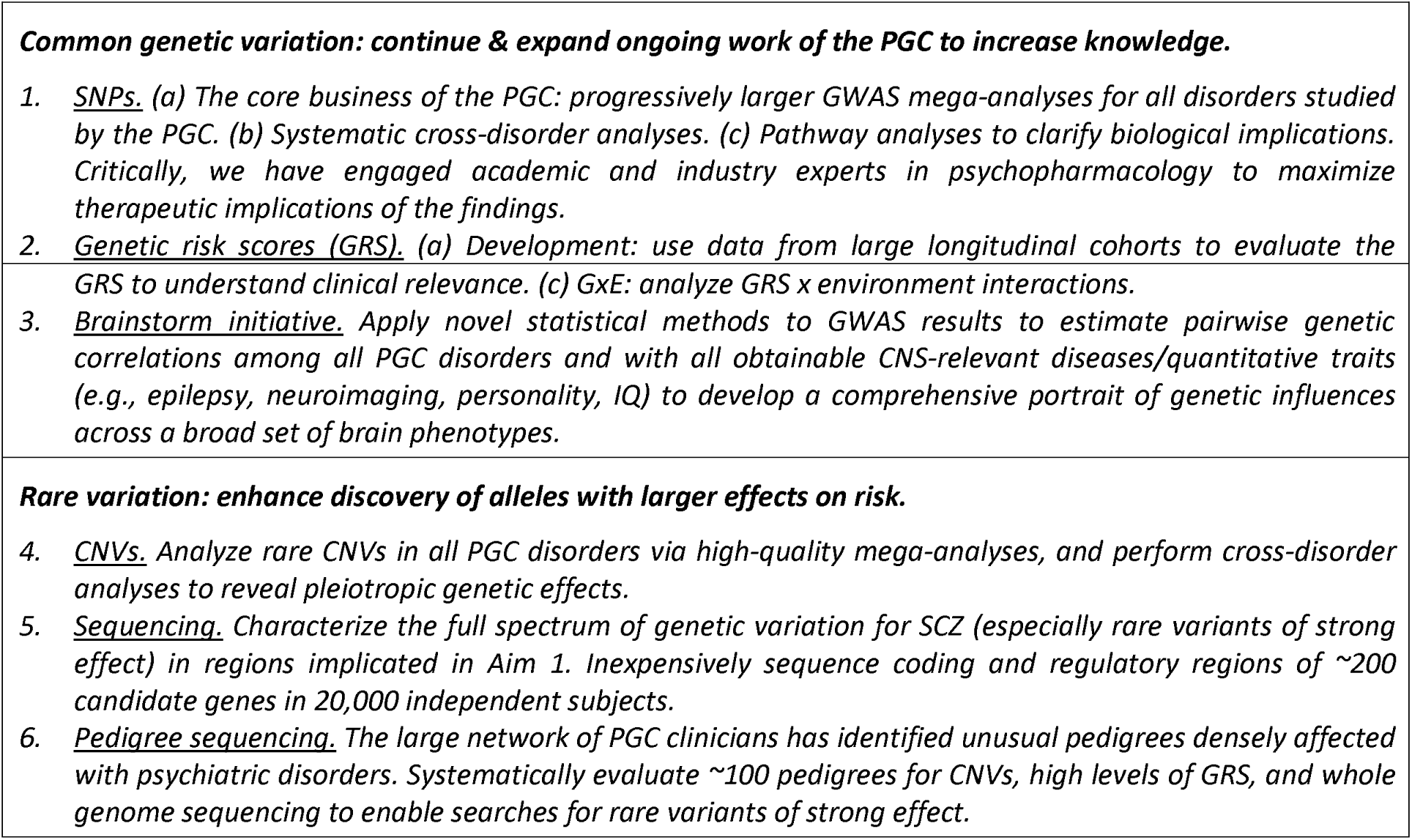
Current projects for the PGC.

### Aim 1

(common variation) is the core business of the PGC: to conduct progressively larger GWAS mega-analyses and systematic cross-disorder analyses (36). *Figure 3a* depicts the progression of sample sizes with time. Our goal is for each of the nine disorder working groups to obtain GWAS data on 100,000 cases. More information on case definitions can be found in *Table S3.*

**Figure 3a.**
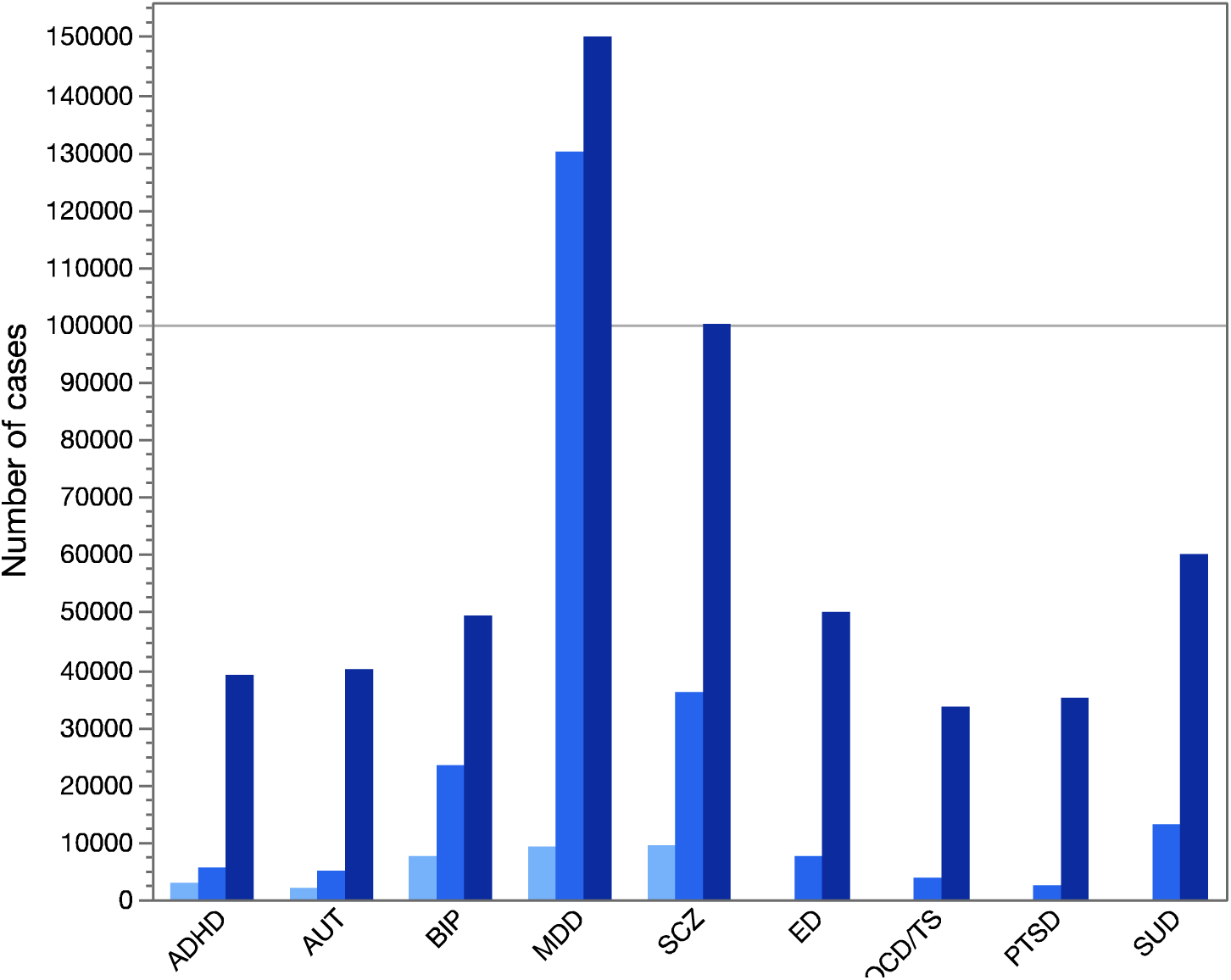
Numbers of cases for PGC GWAS analyses. For the original five PGC disorders (ADHD..SCZ), there are three bars for the numbers of cases in the initial “PGC1” reports, the next round of papers (“PGC2”), and the projected numbers by 2019. For the four disorders added in 2013 (ED..SUD), the PGC2 and projected PGC3 numbers are shown. Abbreviations: ADHD=attention-deficit hyperactivity disorder, AUT=autism, BIP=bipolar disorder, MDD=major depressive disorder, SCZ=schizophrenia, ED=eating disorders, OCD/TS=obsessive-compulsive disorder/Tourette syndrome, PTSD=post-traumatic stress disorder, and SUD=substance use disorders.

*Figure 3b* encapsulates experience with sample size and numbers of significant associations. Some disorders have a fortuitous architecture; e.g., inflammatory bowel disease obtained a considerable number of findings with relatively small samples. For most other complex traits, the path is slower but, with sufficient samples, discovery becomes linear. *Figure 3c* shows an idealized cartoon of the sigmoid-like discovery process from “dead zone” to asymptote. We suggest that the goal is to get to a “good enough” point where most genes are identified at least once and the majority of genes in salient biological processes are highlighted. This can provide an etiologic scaffold for studies that use other methods to identify interacting partners in gene networks and pathways that underlie pathogenesis. There may be on the order of 1,000 genes involved in schizophrenia (37) (for comparison, ^~^13,000 genes are expressed in brain and ^~^2,000 at the synapse). Most PGC groups have at least one association, several are accumulating moderate numbers of loci, and schizophrenia and MDD appear to be in the linear phase (*Table 1b*).

**Figure 3b.**
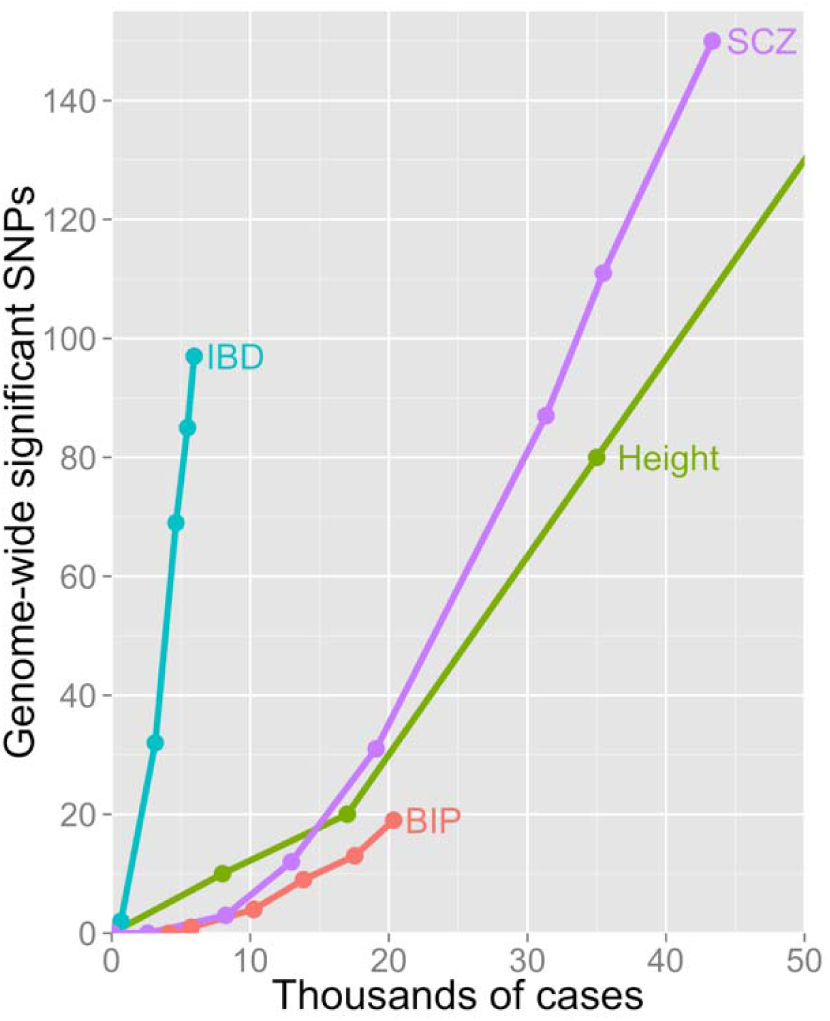
Relation between numbers of cases and genome-wide significant SNPs in GWAS. Lines show discovery paths for inflammatory bowel disease (IBD), schizophrenia (SCZ), height, and bipolar disorder (BIP). IBD has an exceptional genetic architecture and excellent clinical diagnostic specificity that enabled considerable discovery with relatively smaller numbers of cases. SCZ, height, and BIP follow more typical and approximately similar discover paths

**Figure 3c.**
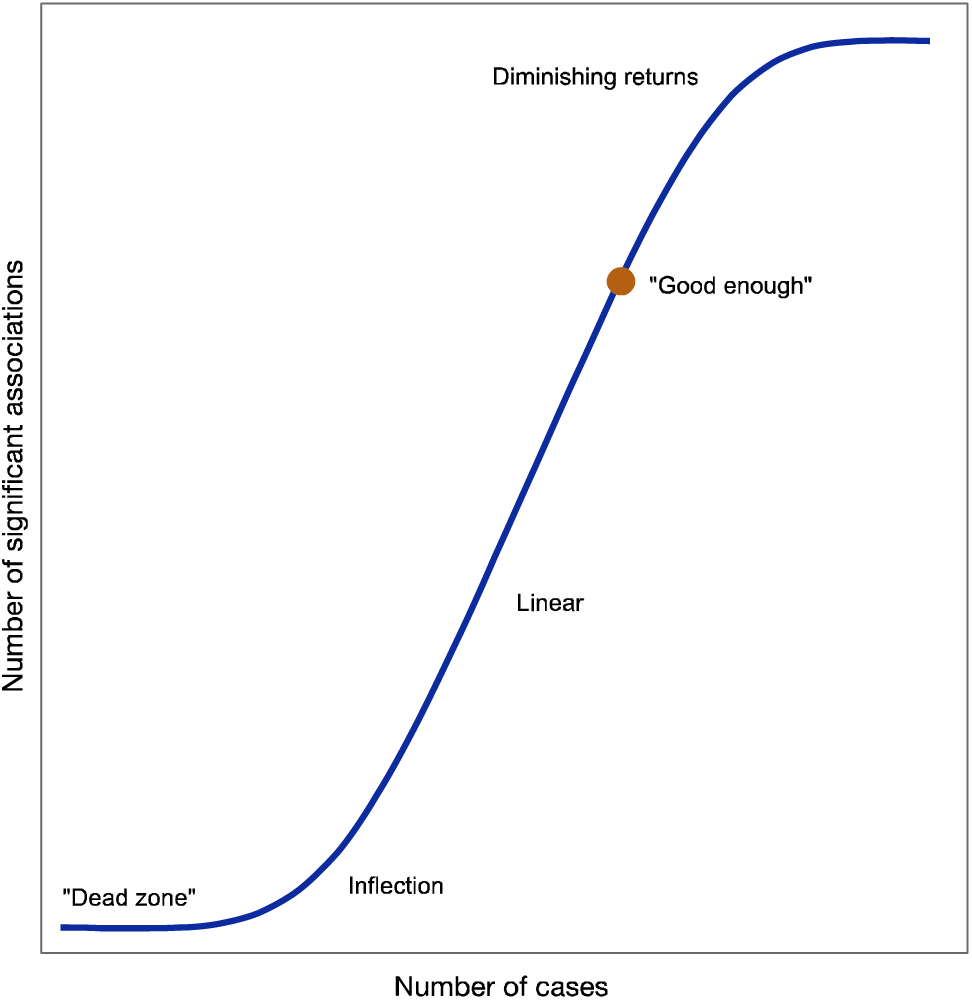
Hypothetical cartoon of the relation between number of cases and genome-wide significant associations for a human complex disease or trait. Complexities arising from the true nature of the initially unknown genetic architecture could change the form of this curve importantly. There is an initial dead zone whose length depends how many cases are accrued and the largest effect size. This is followed by an inflection point where the significant associations begin to accumulate and then a linear phase.

The PGC has extended its initial efforts in three ways. First, we added four new and highly motivated groups (eating disorders, obsessive-compulsive disorder/Tourette syndrome, post-traumatic stress disorder, and substance use disorders). Provisional groups for anxiety disorders and Alzheimer’s disease have been formed. Second, we hope to markedly increase inclusion of non-European samples (*Figure S1*). For example, the PGC is now completing a paper based on over 12,000 schizophrenia cases from East Asia. The post-traumatic stress disorder and substance use disorders groups are studying increasingly larger samples of African-Americans. The Stanley Center of the Broad Institute has launched major sample collection efforts for multiple severe psychiatric disorders in Africa, South America, and Asia.

This work is crucial for generalizability, and it is likely that most (but not all) associations will be observed across the world. Finally, the PGC has engaged academic and industry experts to understand the therapeutic salience of the findings (38). Indeed, the empirical targets of antipsychotic medications are markedly enriched for the results of schizophrenia GWAS and this enrichment became clearer with increasing sample sizes, as has the potential pharmacological relevance of calcium channels for psychiatry (39). The design of rational therapeutics has been an elusive goal for psychiatric indications, and improved genomic knowledge is a pre-competitive activity that can make novel drug discovery more efficient (40).

### Aim 2

(common variation) concerns the analysis of genetic risk scores (GRS). For a complex disease or trait, GRS is a single, normally distributed variable that captures the cumulative effect of risk alleles inherited by an individual (e.g., for schizophrenia, bipolar disorder, or body mass index). Computing a GRS requires a training set (i.e., GWAS results) and genome-wide genotypes on independent test subjects (e.g., a population cohort or participants in a clinical trial). The PGC has made training sets publicly available for multiple disorders (URLs). This allows researchers to compute GRS for whatever use they deem appropriate. GRS are not yet sufficiently discriminating to be useful clinically (14) but are among the first demonstrably valid biomarkers for psychiatric disorders. GRS derived from PGC results have been widely used in psychiatric research for generating patient strata, exploring diagnostic boundaries, identifying cognitive and behavioral correlates of genetic risk that predate clinical disorders, and evaluating the validity of putative cognitive or imaging phenotypes (41). Even social scientists have embraced the approach, seeing opportunities to study how genetic factors interact with the social environment to influence health and broader outcomes (42).

The PGC will systematically evaluate GRS in three contexts: (a) Development: use data from large longitudinal cohorts to evaluate the developmental effects of GRS; (b) Clinical symptoms: analyze relationship between clinical descriptors and GRS to understand clinical relevance; and (c) analyze GRS by environment interactions in epidemiological samples.

### Aim 3

(common variation) will use GWAS results to estimate pairwise genetic correlations among all PGC disorders with all obtainable CNS-relevant diseases and quantitative traits (e.g., epilepsy, neuroimaging, personality, IQ). We will develop a comprehensive portrait of genetic influences across a broad set of brain phenotypes with the intention of improving nosology.

For example, the pairwise genetic correlations (URLs) among anorexia nervosa, ADHD, ASD, bipolar disorder, MDD, and schizophrenia are almost all significant and positive (the exceptions are all correlations of ASD and anorexia nervosa-ADHD, certainly influenced by small sample sizes) (43). This suggests that the common variant genetic architectures – the fundamental liability to these disorders - overlap importantly. After evaluating the disorder-level genetic correlations (22), we will systematically expand our queries to encompass within disorder genetic correlations (e.g., male vs female, early vs later onset) as well as genetic correlations with putative components (e.g., cognition, personality, body mass).

### Aim 4

(rare variation) will continue the PGC’s CNV efforts (29). The PGC CNV group has created a pipeline to call CNVs from the initial intensity files using multiple algorithms followed by careful quality control and analysis. The initial schizophrenia paper has been published, and this group is now working on bipolar disorder, ADHD, and PTSD, and will include more groups with time.

### Aim 5

(rare variation) is a “cheap-seq” aim. We will conduct inexpensive (^~^$50/subject) schizophrenia-focused sequencing of 200 genes in 20,000 subjects. Genes will be selected based on all available sequencing results. For 200 genes, we will increase power far more cost-effectively than whole exome (l0× cheaper) or whole genome sequencing (25× cheaper) in the same time frame. We propose an efficient and affordable way to markedly increase sample sizes for the most promising loci in a new sample of 20,000 subjects.

### Aim 6

(rare variation) will systematically evaluate ^~^100 large pedigrees to search for genetic variants of large effect. We have engaged the large network of PGC clinicians in this task. Most experienced clinicians have encountered unusual pedigrees with high concentrations of severe psychiatric disorders. For example, one pedigree has >100 individuals with a severe psychiatric disorder and eight pedigrees have ≥20 affected individuals. Other pedigrees are from genetic isolates where inter-pedigree marriage is common. Still other pedigrees have extensive comorbidity with intellectual disability and epilepsy. No one has systematically and comprehensively evaluated a large collection of densely affected pedigrees using comprehensive genomic assays (karyotyping, identity-by-descent, CNVs, whole genome sequencing, and GRS) combined with a rigorous statistical framework.

### Actionability

For the common variant aims, Aim 1 is of biological, clinical, and therapeutic relevance. Aim 2 and Aim 3 are important clinically and for nosology. Of the rare variant aims, all are important biologically and therapeutically (given their potential to identify single genes whose mutational disruption carries high risk).

## Issues in the process of being solved

Empirical results from psychiatric genomics have begun to answer many questions. However, we point to two major unresolved issues. First, many GWAS signals are complex. Exactly how these connect to specific genes to enable precise experimentation is solvable but requires functional genomic or functional cellular data. *Figure 4a* shows the *CACNA1C* intronic association for schizophrenia; a subsequent study suggested that these SNPs interact with a regulatory element for *CACNA1C* (44). *Figure 4b* depicts the region surrounding *DRD2* (which encodes the protein targeted by most antipsychotics). This association has been functionally connected to the gene via DNA-DNA interactions (45). *Figure 4c* shows a multigenic region – the association region covers many genes (most of which are expressed in brain) that have been associated with multiple human traits. Finally, *Figure 4d* depicts an intergenic region that is associated with schizophrenia but not near a known protein-coding gene.

**Figure 4.**
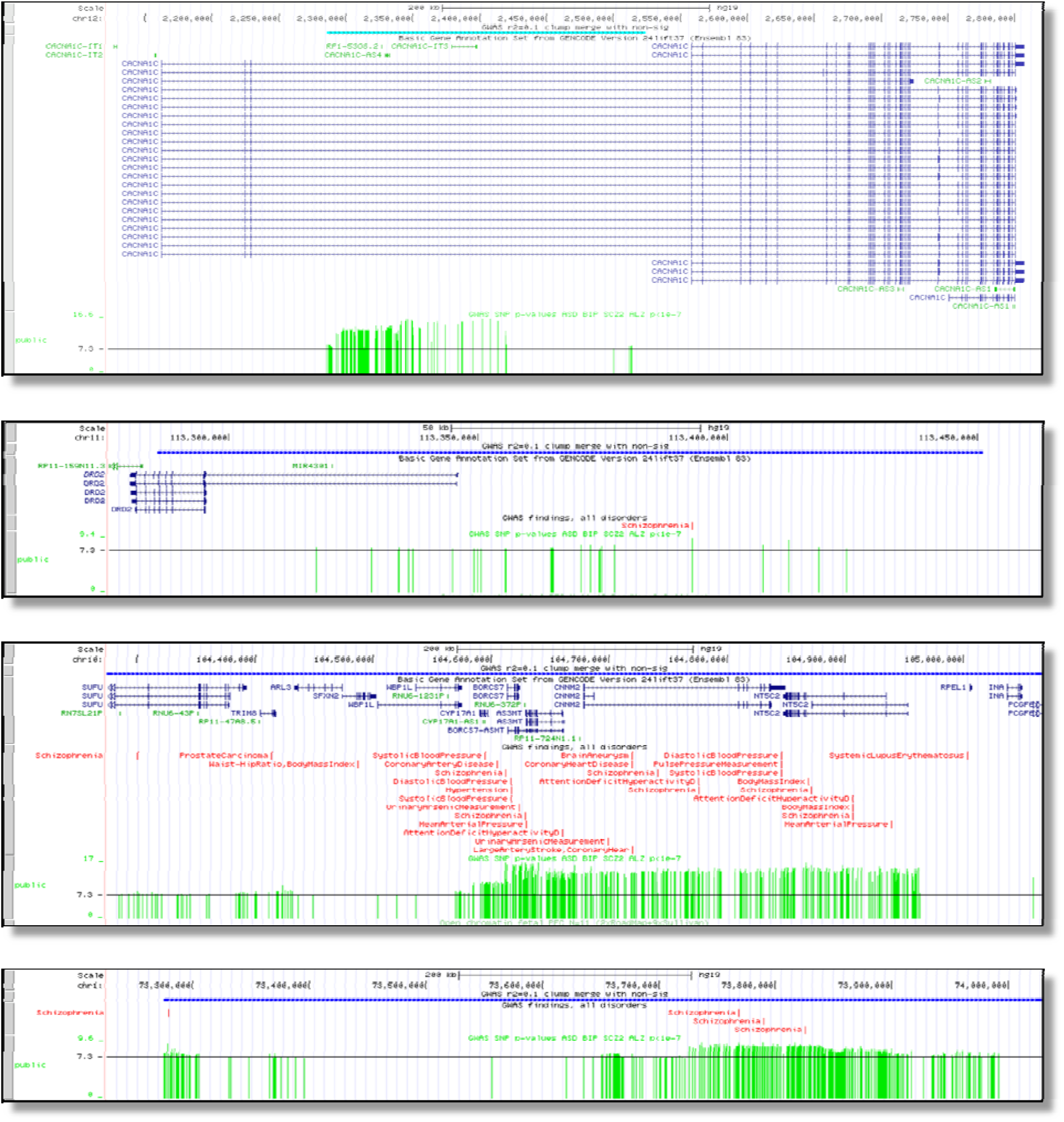
Examples of genome-wide significant regions for schizophrenia with tracks showing the location (hgl9), any genes in the region, GWAS results from the literature, and the schizophrenia results (one green vertical bar per SNP, height corresponds to −log_10_(P-value} with 7.3 equivalent to 5×10^−8^. (a) Intronic association in CACNA1C, (b) association mostly upstream of DRD2, (c) multigenic association, and (d) intergenic association.

These are typical for GWAS results, identifying relatively broad loci with multiple genome-wide significant markers. Although localization is imprecise, the associated genomic regions are clearly informative as they implicate salient biological pathways (46) and increasingly specific genomic features (47). Full elucidation requires functional genomic data generated from relevant cell types including gene expression (48), DNA-DNA looping (45), and epigenomics (49). To facilitate this work, the NIMH has funded the psychENCODE consortium (49) to conduct an array of functional genomic assays on brain tissue from people with severe psychiatric disorders and controls (URLs).

Second, the extant cross-disorder studies of psychiatric disorders show non-trivial genetic overlap between most or all disorders. The results clearly show that our diagnostic categories do not define pathophysiological entities. It is plausible that a focus on components would be valuable (e.g., cognition, mood stability, or innate anxiety-fearfulness). The resolution of these issues will address major unanswered questions: from a genetic perspective, what are these disorders? How are they similar and how are they different?

## Common complaints

Briefly, there are three common complaints about the work of the PGC. First, “the results don’t matter” – the readouts are broad with small effect sizes, and the effect sizes of individual associated loci are small. In fact, the results are delivering increasingly useful and targeted knowledge (discussed under Aim 1 above) (38, 39, 46). The small effect sizes do not constrain the potential utility of targeting the identified genes or pathways – drugs targeting those pathways can have major effects. Unlike 10 years ago, pharmaceutical companies are following this area closely as genomic data are increasingly crucial to drug development (40).

Second, “what about unaccounted heritability (h^2^)?” Heritability estimated from genome-wide SNP data (SNP-h^2^) depends on technical issues and sample size. The comparator is estimated from twin or family data (twin-h^2^ or pedigree-h^2^), but these are assumption-laden and imprecise. “Unaccounted h^2^” is the difference between these, and attempts to reconcile two fundamentally different entities. Still, when the genomic study is sufficiently large as with schizophrenia, SNP-h^2^ is around half of the pedigree-h^2^. A point often missed, however, is that explaining h^2^ is a minor goal. The main goals of the PGC are to gain biological, clinical, and therapeutic insights, which can arise regardless of the magnitude of heritability accounted for.

Third, because most PGC analyses are based on categorical, case vs control analyses, “PGC cases lack clinical depth”. This was by intention: over 10 years ago (37), some of us reasoned that fast phenotype characterization that led to affordably large sample sizes was the logical first step (as opposed to large numbers of phenotypes on small numbers of subjects). This was always the first step. The success of this strategy is seen, not only in the genome-wide significant loci that we have discovered, but also in the many phenotypes that have been associated with PGC GRS in both clinical and population samples. The second step, ongoing now, is detailed characterization of genetically informative subsets of cases (e.g., Aim 2). In addition, some PGC working groups (e.g., Substance Use Disorders) are currently analyzing quantitative phenotypes.

## Conclusion

The PGC is the largest and most systematic genomics effort in the history of psychiatry. In the next five years, we propose to markedly up-scale our work. By tackling nature as it is and not as we might want it to be, we hope to provide considerable new knowledge about the fundamental basis of psychiatric disorders. Our long-standing commitment to global collaboration, open science, and rapid progress means that we will make our results and tools available in a timely manner. Prediction of the future is always hazardous but, given that we finally have a minimally adequate toolkit for genomics, it is possible that we are entering a golden age of research into the fundamental basis of severe mental illness.

## Conflicts of Interest

PFS is a scientific advisor for Pfizer, Inc. and received an honorarium from F. Hoffmann-La Roche AG. CMB is a consultant for and grant recipient from Shire. Cardiff University received an honorarium from F. Hoffmann-La Roche AG for a presentation by MOD in 2015. OAA received a speaker’s honorarium from Lundbeck in 2016. Multiple drug companies work with the PGC in a manner equivalent to academic investigators.

## Acknowledgements

We are deeply indebted to the investigators who comprise the PGC, and to the hundreds of thousands of subjects who have shared their life experiences with PGC investigators. The PGC has received major funding from the US National Institute of Mental Health and the US National Institute of Drug Abuse (PGC3: U01 MH109528 and U01 MH109532, PGC2: U01 MH094421, PGC1: U01 MH085520). AA also acknowledges K02 DA32573. Other significant funders include the Lundbeck Foundation, Stanley Center of the Broad Institute, Science Foundation Ireland, Cohen Veterans Bioscience, and the Norwegian Institute of Health. SVF is supported by the K.G. Jebsen Centre for Research on Neuropsychiatric Disorders, University of Bergen, Bergen, Norway, the European Union’s Seventh Framework Programme for research, technological development and demonstration under grant agreement no 602805, the European Union’s Horizon 2020 research and innovation programme under grant agreement No 667302 and NIMH grants 5R01MH101519 and U01 MH109536-01.

## URLs

European Bioinformatics Institute, https://www.ebi.ac.uk/Rwas

LD-Hub, http://ldsc.broadinstitute.org

psychENCODE, https://psychencode.org

Psychiatric Genomics Consortium, general information, https://pgc.unc.edu

Psychiatric Genomics Consortium, results, http://www.med.unc.edu/pgc/results-and-downloads

Psychiatric Genomics Consortium, monthly talks, http://www.med.unc.edu/pgc/worldwide

## References

1. Kendler KS, Prescott CA: Genes, Environment, and Psychopathology. New York, Guilford Press; 2006.

2. Polderman TJ, Benyamin B, de Leeuw CA, Sullivan PF, van Bochoven A, Visscher PM, Posthuma D. Fifty years of twin studies: A meta-analysis of the heritability of human traits. Nature Genetics. 2015;47702–709.

3. Yang J, Lee SH, Goddard ME, Visscher PM. GCTA: a tool for genome-wide complex trait analysis. American journal of human genetics. 2011;88:76–82.

4. Bulik-Sullivan BK, Finucane HK, Anttila V, Gusev A, Day FR, ReproGen Consortium, Psychiatric Genomics Consortium, Anorexia Nervosa Genetic Consortium, Duncan L, Perry JRB, Patterson N, Robinson E, Daly MJ, Price AL, Neale BM. An atlas of genetic correlations across human diseases and traits. Nature Genetics. 2015;47:1236–1241.

5. Psychiatric GWAS Consortium. A framework for interpreting genome-wide association studies of psychiatric disorders. Molecular Psychiatry. 2009;14:10–17.

6. Psychiatric GWAS Consortium. Genome-wide association studies: history, rationale, and prospects for psychiatric disorders. Am J Psychiatry. 2009;166:540–546.

7. Cross Disorder Phenotype Group of the Psychiatric GWAS Consortium. Dissecting the phenotype in genome-wide association studies of psychiatric illness. Br J Psychiatry. 2009;195:97–99.

8. Lambert JC, Ibrahim-Verbaas CA, Harold D, Naj AC, Sims R, Bellenguez C, DeStafano AL, Bis JC, Beecham GW, Grenier-Boley B, Russo G, Thorton-Wells TA, Jones N, Smith AV, Chouraki V, Thomas C, Ikram MA, Zelenika D, Vardarajan BN, Kamatani Y, Lin CF, Gerrish A, Schmidt H, Kunkle B, Dunstan ML, Ruiz A, Bihoreau MT, Choi SH, Reitz C, Pasquier F, Cruchaga C, Craig D, Amin N, Berr C, Lopez OL, De Jager PL, Deramecourt V, Johnston JA, Evans D, Lovestone S, Letenneur L, Moron FJ, Rubinsztein DC, Eiriksdottir G, Sleegers K, Goate AM, Fievet N, Huentelman MW, Gill M, Brown K, Kamboh MI, Keller L, Barberger-Gateau P, McGuiness B, Larson EB, Green R, Myers AJ, Dufouil C, Todd S, Wallon D, Love S, Rogaeva E, Gallacher J, St George-Hyslop P, Clarimon J, Lleo A, Bayer A, Tsuang DW, Yu L, Tsolaki M, Bossu P, Spalletta G, Proitsi P, Collinge J, Sorbi S, Sanchez-Garcia F, Fox NC, Hardy J, Deniz Naranjo MC, Bosco P, Clarke R, Brayne C, Galimberti D, Mancuso M, Matthews F, European Alzheimer’s Disease I, Genetic, Environmental Risk in Alzheimer’s D, Alzheimer’s Disease Genetic C, Cohorts for H, Aging Research in Genomic E, Moebus S, Mecocci P, Del Zompo M, Maier W, Hampel H, Pilotto A, Bullido M, Panza F, Caffarra P, Nacmias B, Gilbert JR, Mayhaus M, Lannefelt L, Hakonarson H, Pichler S, Carrasquillo MM, Ingelsson M, Beekly D, Alvarez V, Zou F, Valladares O, Younkin SG, Coto E, Hamilton-Nelson KL, Gu W, Razquin C, Pastor P, Mateo I, Owen MJ, Faber KM, Jonsson PV, Combarros O, O’Donovan MC, Cantwell LB, Soininen H, Blacker D, Mead S, Mosley TH Jr., Bennett DA, Harris TB, Fratiglioni L, Holmes C, de Bruijn RF, Passmore P, Montine TJ, Bettens K, Rotter JI, Brice A, Morgan K, Foroud TM, Kukull WA, Hannequin D, Powell JF, Nalls MA, Ritchie K, Lunetta KL, Kauwe JS, Boerwinkle E, Riemenschneider M, Boada M, Hiltuenen M, Martin ER, Schmidt R, Rujescu D, Wang LS, Dartigues JF, Mayeux R, Tzourio C, Hofman A, Nothen MM, Graff C, Psaty BM, Jones L, Haines JL, Holmans PA, Lathrop M, Pericak-Vance MA, Launer LJ, Farrer LA, van Duijn CM, Van Broeckhoven C, Moskvina V, Seshadri S, Williams J, Schellenberg GD, Amouyel P. Meta-analysis of 74,046 individuals identifies 11 new susceptibility loci for Alzheimer’s disease. Nat Genet. 2013;45:1452–1458.

9. Klein RJ, Zeiss C, Chew EY, Tsai JY, Sackler RS, Haynes C, Henning AK, Sangiovanni JP, Mane SM, Mayne ST, Bracken MB, Ferris FL, Ott J, Barnstable C, Hoh J. Complement factor H polymorphism in age-related macular degeneration. Science. 2005;308:385–389.

10. International Schizophrenia Consortium. Common polygenic variation contributes to risk of schizophrenia and bipolar disorder. Nature. 2009;460:748–752.

11. WTCCC. Genome-wide association study of 14,000 cases of seven common diseases and 3,000 shared controls. Nature. 2007;447:661–678.

12. Wray N, Pergadia M, Blackwood D, Penninx B, Gordon S, Nyholt D, Ripke S, MacIntyre D, McGhee K, Maclean A, Smit J, Hottenga J, Willemsen G, Middeldorp C, de Geus E, Lewis C, McGuffin P, Hickie I, van den Oord E, Liu J, Macgregor S, McEvoy B, Byrne E, Medland S, Statham D, Henders A, Heath A, Montgomery G, Martin N, Boomsma D, Madden P, Sullivan PF. Genomewide association study of major depressive disorder: New results, meta-analysis, and lessons learned. Molecular Psychiatry. 2011;17:36–48.

13. Neale BM, Medland SE, Ripke S, Asherson P, Franke B, Lesch KP, Faraone SV, Nguyen TT, Schafer H, Holmans P, Daly M, Steinhausen HC, Freitag C, Reif A, Renner TJ, Romanos M, Romanos J, Walitza S, Warnke A, Meyer J, Palmason H, Buitelaar J, Vasquez AA, Lambregts-Rommelse N, Gill M, Anney RJ, Langely K, O’Donovan M, Williams N, Owen M, Thapar A, Kent L, Sergeant J, Roeyers H, Mick E, Biederman J, Doyle A, Smalley S, Loo S, Hakonarson H, Elia J, Todorov A, Miranda A, Mulas F, Ebstein RP, Rothenberger A, Banaschewski T, Oades RD, Sonuga-Barke E, McGough J, Nisenbaum L, Middleton F, Hu X, Nelson S, ADHD Working Group of the Psychiatric GWAS. Consortium. Meta-analysis of genome-wide association studies of attention-deficit/hyperactivity disorder. J Am Acad Child Adolesc Psychiatry. 2010;49:884–897.

14. Schizophrenia Working Group of the Psychiatric Genomics Consortium. Biological insights from 108 schizophrenia-associated genetic loci. Nature. 2014;511:421–427.

15. Takeuchi F, Isono M, Nabika T, Katsuya T, Sugiyama T, Yamaguchi S, Kobayashi S, Ogihara T, Yamori Y, Fujioka A, Kato N. Confirmation of ALDH2 as a Major locus of drinking behavior and of its variants regulating multiple metabolic phenotypes in a Japanese population. Circ J. 2011;75:911–918.

16. Goldstein JI, Jarskog LF, Hilliard C, Alfirevic A, Duncan LE, Fourches D, Huang H, Lehner T, Lek M, Neale BM, Ripke S, Shianna KV, Szatkiewicz JP, Tropsha A, van den Oord EJCG, Cascorbi I, Dettling M, Gazit E, Goff DC, Holden AL, Kelly DL, Malhotra AK, Nielsen J, Pirmohamed M, Rujescu D, Werge T, Levy DL, Josiassen RC, Kennedy JL, Lieberman JA, Daly MJ, Sullivan PF. Clozapine-induced agranulocytosis/granulocytopenia is associated with rare *HLA-DQB1* and *HLA-B* alleles. Nature communications. 2014;5:4757.

17. McClellan JM, Susser E, King MC. Schizophrenia: a common disease caused by multiple rare alleles. Br J Psychiatry. 2007;190:194–199.

18. Goldstein DB. Common genetic variation and human traits. N Engl J Med. 2009;360:1696–1698.

19. Malhotra D, Sebat J. CNVs: Harbingers of a Rare Variant Revolution in Psychiatric Genetics. Cell. 2012;148:1223–1241.

20. Sullivan PF, Psychiatric Genomics Consortium. Don’t give up on GWAS. Molecular Psychiatry. 2011;17:2–3.

21. Cross-Disorder Group of the Psychiatric Genomics Consortium. Genetic relationship between five psychiatric disorders estimated from genome-wide SNPs. Nature genetics. 2013;45:984–994.

22. Anttila V, Bulik-Sullivan BK, Finucane HK, Bras J, Duncan L, Escott-Price V, Falcone G, Gormley P, Malik R, Patsopoulos N, Ripke S, Walters R, Wei Z, Yu D, Lee P, IGAP consortium, IHGC consortium, ILAE Consortium on Complex Epilepsies, IMSGC consortium, IPDGC consortium, METASTROKE and ICH Studies of the ISGC, ADHD Working Group of the PGC, Anorexia Nervosa Working Group of the PGC, ASD Working Group of the PGC, Bipolar Disorders Working Group of the PGC, Major Depressive Disorder Working Group of the PGC, OCD and TS Working Group of the PGC, Schizophrenia Working Group of the PGC, Breen G, Bulik C, Daly M, Dichgans M, Faraone S, Guerreiro R, Holmans P, Kendler K, Koeleman B, Mathews C, Scharf J, Sklar P, Williams J, Wood N, Cotsapas C, Palotie A, Smoller J, Sullivan PF, Rosand J, Corvin A, Neale B. Analysis of shared heritability in common disorders of the brain. Submitted

23. Gaugler T, Klei L, Sanders SJ, Bodea CA, Goldberg AP, Lee AB, Mahajan M, Manaa D, Pawitan Y, Reichert J, Ripke S, Sandin S, Sklar P, Svantesson O, Reichenberg A, Hultman CM, Devlin B, Roeder K, Buxbaum JD. Most genetic risk for autism resides with common variation. Nat Genet. 2014.

24. Fuchsberger C, Flannick J, Teslovich TM, Mahajan A, Agarwala V, Gaulton KJ, Ma C, Fontanillas P, Moutsianas L, McCarthy DJ, Rivas MA, Perry JR, Sim X, Blackwell TW, Robertson NR, Rayner NW, Cingolani P, Locke AE, Fernandez Tajes J, Highland HM, Dupuis J, Chines PS, Lindgren CM, Hartl C, Jackson AU, Chen H, Huyghe JR, van de Bunt M, Pearson RD, Kumar A, Muller-Nurasyid M, Grarup N, Stringham HM, Gamazon ER, Lee J, Chen Y, Scott RA, Below JE, Chen P, Huang J, Go MJ, Stitzel ML, Pasko D, Parker SC, Varga TV, Green T, Beer NL, Day-Williams AG, Ferreira T, Fingerlin T, Horikoshi M, Hu C, Huh I, Ikram MK, Kim BJ, Kim Y, Kim YJ, Kwon MS, Lee J, Lee S, Lin KH, Maxwell TJ, Nagai Y, Wang X, Welch RP, Yoon J, Zhang W, Barzilai N, Voight BF, Han BG, Jenkinson CP, Kuulasmaa T, Kuusisto J, Manning A, Ng MC, Palmer ND, Balkau B, Stancakova A, Abboud HE, Boeing H, Giedraitis V, Prabhakaran D, Gottesman O, Scott J, Carey J, Kwan P, Grant G, Smith JD, Neale BM, Purcell S, Butterworth AS, Howson JM, Lee HM, Lu Y, Kwak SH, Zhao W, Danesh J, Lam VK, Park KS, Saleheen D, So WY, Tam CH, Afzal U, Aguilar D, Arya R, Aung T, Chan E, Navarro C, Cheng CY, Palli D, Correa A, Curran JE, Rybin D, Farook VS, Fowler SP, Freedman BI, Griswold M, Hale DE, Hicks PJ, Khor CC, Kumar S, Lehne B, Thuillier D, Lim WY, Liu J, van der Schouw YT, Loh M, Musani SK, Puppala S, Scott WR, Yengo L, Tan ST, Taylor HA, Jr., Thameem F, Wilson G, Sr., Wong TY, Njolstad PR, Levy JC, Mangino M, Bonnycastle LL, Schwarzmayr T, Fadista J, Surdulescu GL, Herder C, Groves CJ, Wieland T, Bork-Jensen J, Brandslund I, Christensen C, Koistinen HA, Doney AS, Kinnunen L, Esko T, Farmer AJ, Hakaste L, Hodgkiss D, Kravic J, Lyssenko V, Hollensted M, Jorgensen ME, Jorgensen T, Ladenvall C, Justesen JM, Karajamaki A, Kriebel J, Rathmann W, Lannfelt L, Lauritzen T, Narisu N, Linneberg A, Melander O, Milani L, Neville M, Orho-Melander M, Qi L, Qi Q, Roden M, Rolandsson O, Swift A, Rosengren AH, Stirrups K, Wood AR, Mihailov E, Blancher C, Carneiro MO, Maguire J, Poplin R, Shakir K, Fennell T, DePristo M, Hrabe de Angelis M, Deloukas P, Gjesing AP, Jun G, Nilsson P, Murphy J, Onofrio R, Thorand B, Hansen T, Meisinger C, Hu FB, Isomaa B, Karpe F, Liang L, Peters A, Huth C, O’Rahilly SP, Palmer CN, Pedersen O, Rauramaa R, Tuomilehto J, Salomaa V, Watanabe RM, Syvanen AC, Bergman RN, Bharadwaj D, Bottinger EP, Cho YS, Chandak GR, Chan JC, Chia KS, Daly MJ, Ebrahim SB, Langenberg C, Elliott P, Jablonski KA, Lehman DM, Jia W, Ma RC, Pollin TI, Sandhu M, Tandon N, Froguel P, Barroso I, Teo YY, Zeggini E, Loos RJ, Small KS, Ried JS, DeFronzo RA, Grallert H, Glaser B, Metspalu A, Wareham NJ, Walker M, Banks E, Gieger C, Ingelsson E, Im HK, Illig T, Franks PW, Buck G, Trakalo J, Buck D, Prokopenko I, Magi R, Lind L, Farjoun Y, Owen KR, Gloyn AL, Strauch K, Tuomi T, Kooner JS, Lee JY, Park T, Donnelly P, Morris AD, Hattersley AT, Bowden DW, Collins FS, Atzmon G, Chambers JC, Spector TD, Laakso M, Strom TM, Bell GI, Blangero J, Duggirala R, Tai ES, McVean G, Hanis CL, Wilson JG, Seielstad M, Frayling TM, Meigs JB, Cox NJ, Sladek R, Lander ES, Gabriel S, Burtt NP, Mohlke KL, Meitinger T, Groop L, Abecasis G, Florez JC, Scott LJ, Morris AP, Kang HM, Boehnke M, Altshuler D, McCarthy ML. The genetic architecture of type 2 diabetes. Nature. 2016;536:41–47.

25. Wood AR, Esko T, Yang J, Vedantam S, Pers TH, Gustafsson S, Chu AY, Estrada K, Luan J, Kutalik Z, Amin N, Buchkovich ML, Croteau-Chonka DC, Day FR, Duan Y, Fall T, Fehrmann R, Ferreira T, Jackson AU, Karjalainen J, Lo KS, Locke AE, Magi R, Mihailov E, Porcu E, Randall JC, Scherag A, Vinkhuyzen AA, Westra HJ, Winkler TW, Workalemahu T, Zhao JH, Absher D, Albrecht E, Anderson D, Baron J, Beekman M, Demirkan A, Ehret GB, Feenstra B, Feitosa MF, Fischer K, Fraser RM, Goel A, Gong J, Justice AE, Kanoni S, Kleber ME, Kristiansson K, Lim U, Lotay V, Lui JC, Mangino M, Mateo Leach I, Medina-Gomez C, Nalls MA, Nyholt DR, Palmer CD, Pasko D, Pechlivanis S, Prokopenko I, Ried JS, Ripke S, Shungin D, Stancakova A, Strawbridge RJ, Sung YJ, Tanaka T, Teumer A, Trompet S, van der Laan SW, van Setten J, Van Vliet-Ostaptchouk JV, Wang Z, Yengo L, Zhang W, Afzal U, Arnlov J, Arscott GM, Bandinelli S, Barrett A, Bellis C, Bennett AJ, Berne C, Bluher M, Bolton JL, Bottcher Y, Boyd HA, Bruinenberg M, Buckley BM, Buyske S, Caspersen IH, Chines PS, Clarke R, Claudi-Boehm S, Cooper M, Daw EW, De Jong PA, Deelen J, Delgado G, Denny JC, Dhonukshe-Rutten R, Dimitriou M, Doney AS, Dorr M, Eklund N, Eury E, Folkersen L, Garcia ME, Geller F, Giedraitis V, Go AS, Grallert H, Grammer TB, Grassler J, Gronberg H, de Groot LC, Groves LC, Haessler J, Hall P, Haller T, Hallmans G, Hannemann A, Hartman CA, Hassinen M, Hayward C, Heard-Costa NL, Helmer Q, Hemani G, Henders AK, Hillege HL, Hlatky MA, Hoffmann W, Hoffmann P, Holmen O, Houwing-Duistermaat JJ, Illig T, Isaacs A, James AL, Jeff J, Johansen B, Johansson A, Jolley J, Juliusdottir T, Junttila J, Kho AN, Kinnunen L, Klopp N, Kocher T, Kratzer W, Lichtner P, Lind L, Lindstrom J, Lobbens S, Lorentzon M, Lu Y, Lyssenko V, Magnusson PK, Mahajan A, Maillard M, McArdle WL, McKenzie CA, McLachlan S, McLaren PJ, Menni C, Merger S, Milani L, Moayyeri A, Monda KL, Morken MA, Muller G, Muller-Nurasyid M, Musk AW, Narisu N, Nauck M, Nolte IM, Nothen MM, Oozageer L, Pilz S, Rayner NW, Renstrom F, Robertson NR, Rose LM, Roussel R, Sanna S, Scharnagl H, Scholtens S, Schumacher FR, Schunkert H, Scott RA, Sehmi J, Seufferlein T, Shi J, Silventoinen K, Smit JH, Smith AV, Smolonska J, Stanton AV, Stirrups K, Stott DJ, Stringham HM, Sundstrom J, Swertz MA, Syvanen AC, Tayo BO, Thorleifsson G, Tyrer JP, van Dijk S, van Schoor NM, van der Velde N, van Heemst D, van Oort FV, Vermeulen SH, Verweij N, Vonk JM, Waite LL, Waldenberger M, Wennauer R, Wilkens LR, Willenborg C, Wilsgaard T, Wojczynski MK, Wong A, Wright AF, Zhang Q, Arveiler D, Bakker SJ, Beilby J, Bergman RN, Bergmann S, Biffar R, Blangero J, Boomsma DI, Bornstein SR, Bovet P, Brambilla P, Brown MJ, Campbell H, Caulfield MJ, Chakravarti A, Collins R, Collins FS, Crawford DC, Cupples LA, Danesh J, de Faire U, den Ruijter HM, Erbel R, Erdmann J, Eriksson JG, Farrall M, Ferrannini E, Ferrieres J, Ford I, Forouhi NG, Forrester T, Gansevoort RT, Gejman PV, Gieger C, Golay A, Gottesman O, Gudnason V, Gyllensten U, Haas DW, Hall AS, Harris TB, Hattersley AT, Heath AC, Hengstenberg C, Hicks AA, Hindorff LA, Hingorani AD, Hofman A, Hovingh GK, Humphries SE, Hunt SC, Hypponen E, Jacobs KB, Jarvelin MR, Jousilahti P, Jula AM, Kaprio J, Kastelein JJ, Kayser M, Kee F, Keinanen-Kiukaanniemi SM, Kiemeney LA, Kooner JS, Kooperberg C, Koskinen S, Kovacs P, Kraja AT, Kumari M, Kuusisto J, Lakka TA, Langenberg C, Le Marchand L, Lehtimaki T, Lupoli S, Madden PA, Mannisto S, Manunta P, Marette A, Matise TC, McKnight B, Meitinger T, Moll FL, Montgomery GW, Morris AD, Morris AP, Murray JC, Nelis M, Ohlsson C, Oldehinkel AJ, Ong KK, Ouwehand WH, Pasterkamp G, Peters A, Pramstaller PP, Price JF, Qi L, Raitakari OT, Rankinen T, Rao DC, Rice TK, Ritchie M, Rudan I, Salomaa V, Samani NJ, Saramies J, Sarzynski MA, Schwarz PE, Sebert S, Sever P, Shuldiner AR, Sinisalo J, Steinthorsdottir V, Stolk RP, Tardif JC, Tonjes A, Tremblay A, Tremoli E, Virtamo J, Vohl MC, Electronic Medical R, Genomics C, Consortium MI, Consortium P, LifeLines Cohort S, Amouyel P, Asselbergs FW, Assimes TL, Bochud M, Boehm BO, Boerwinkle E, Bottinger EP, Bouchard C, Cauchi S, Chambers JC, Chanock SJ, Cooper RS, de Bakker PI, Dedoussis G, Ferrucci L, Franks PW, Froguel P, Groop LC, Haiman CA, Hamsten A, Hayes MG, Hui J, Hunter DJ, Hveem K, Jukema JW, Kaplan RC, Kivimaki M, Kuh D, Laakso M, Liu Y, Martin NG, Marz W, Melbye M, Moebus S, Munroe PB, Njolstad I, Oostra BA, Palmer CN, Pedersen NL, Perola M, Perusse L, Peters U, Powell JE, Power C, Quertermous T, Rauramaa R, Reinmaa E, Ridker PM, Rivadeneira F, Rotter JI, Saaristo TE, Saleheen D, Schlessinger D, Slagboom PE, Snieder H, Spector TD, Strauch K, Stumvoll M, Tuomilehto J, Uusitupa M, van der Harst P, Volzke H, Walker M, Wareham NJ, Watkins H, Wichmann HE, Wilson JF, Zanen P, Deloukas P, Heid IM, Lindgren CM, Mohlke KL, Speliotes EK, Thorsteinsdottir U, Barroso I, Fox CS, North KE, Strachan DP, Beckmann JS, Berndt SI, Boehnke M, Borecki IB, McCarthy MI, Metspalu A, Stefansson K, Uitterlinden AG, van Duijn CM, Franke L, Willer CJ, Price AL, Lettre G, Loos RJ, Weedon MN, Ingelsson E, O’Connell JR, Abecasis GR, Chasman DI, Goddard ME, Visscher PM, Hirschhorn JN, Frayling TM. Defining the role of common variation in the genomic and biological architecture of adult human height. Nat Genet. 2014;46:1173–1186.

26. Locke AE, Kahali B, Berndt SI, Justice AE, Pers TH, Day FR, Powell C, Vedantam S, Buchkovich ML, Yang J, Croteau-Chonka DC, Esko T, Fall T, Ferreira T, Gustafsson S, Kutalik Z, Luan J, Magi R, Randall JC, Winkler TW, Wood AR, Workalemahu T, Faul JD, Smith JA, Hua Zhao J, Zhao W, Chen J, Fehrmann R, Hedman AK, Karjalainen J, Schmidt EM, Absher D, Amin N, Anderson D, Beekman M, Bolton JL, Bragg-Gresham JL, Buyske S, Demirkan A, Deng G, Ehret GB, Feenstra B, Feitosa MF, Fischer K, Goel A, Gong J, Jackson AU, Kanoni S, Kleber ME, Kristiansson K, Lim U, Lotay V, Mangino M, Mateo Leach I, Medina-Gomez C, Medland SE, Nalls MA, Palmer CD, Pasko D, Pechlivanis S, Peters MJ, Prokopenko I, Shungin D, Stancakova A, Strawbridge RJ, Ju Sung Y, Tanaka T, Teumer A, Trompet S, van der Laan SW, van Setten J, Van Vliet-Ostaptchouk JV, Wang Z, Yengo L, Zhang W, Isaacs A, Albrecht E, Arnlov J, Arscott GM, Attwood AP, Bandinelli S, Barrett A, Bas IN, Bellis C, Bennett AJ, Berne C, Blagieva R, Bluher M, Bohringer S, Bonnycastle LL, Bottcher Y, Boyd HA, Bruinenberg M, Caspersen IH, Ida Chen YD, Clarke R, Daw EW, de Craen AJ, Delgado G, Dimitriou M, Doney AS, Eklund N, Estrada K, Eury E, Folkersen L, Fraser RM, Garcia ME, Geller F, Giedraitis V, Gigante B, Go AS, Golay A, Goodall AH, Gordon SD, Gorski M, Grabe HJ, Grallert H, Grammer TB, Grassler J, Gronberg H, Groves CJ, Gusto G, Haessler J, Hall P, Haller T, Hallmans G, Hartman CA, Hassinen M, Hayward C, Heard-Costa NL, Helmer Q, Hengstenberg C, Holmen O, Hottenga JJ, James AL, Jeff JM, Johansson A, Jolley J, Juliusdottir T, Kinnunen L, Koenig W, Koskenvuo M, Kratzer W, Laitinen J, Lamina C, Leander K, Lee NR, Lichtner P, Lind L, Lindstrom J, Sin Lo K, Lobbens S, Lorbeer R, Lu Y, Mach F, Magnusson PK, Mahajan A, McArdle WL, McLachlan S, Menni C, Merger S, Mihailov E, Milani L, Moayyeri A, Monda KL, Morken MA, Mulas A, Muller G, Muller-Nurasyid M, Musk AW, Nagaraja R, Nothen MM, Nolte IM, Pilz S, Rayner NW, Renstrom F, Rettig R, Ried JS, Ripke S, Robertson NR, Rose LM, Sanna S, Scharnagl H, Scholtens S, Schumacher FR, Scott WR, Seufferlein T, Shi J, Vernon Smith A, Smolonska J, Stanton AV, Steinthorsdottir V, Stirrups K, Stringham HM, Sundstrom J, Swertz MA, Swift AJ, Syvanen AC, Tan ST, Tayo BO, Thorand B, Thorleifsson G, Tyrer JP, Uh HW, Vandenput L, Verhulst FC, Vermeulen SH, Verweij N, Vonk JM, Waite LL, Warren HR, Waterworth D, Weedon MN, Wilkens LR, Willenborg C, Wilsgaard T, Wojczynski MK, Wong A, Wright AF, Zhang Q, LifeLines Cohort S, Brennan EP, Choi M, Dastani Z, Drong AW, Eriksson P, Franco-Cereceda A, Gadin JR, Gharavi AG, Goddard ME, Handsaker RE, Huang J, Karpe F, Kathiresan S, Keildson S, Kiryluk K, Kubo M, Lee JY, Liang L, Lifton RP, Ma B, McCarroll SA, McKnight AJ, Min JL, Moffatt MF, Montgomery GW, Murabito JM, Nicholson G, Nyholt DR, Okada Y, Perry JR, Dorajoo R, Reinmaa E, Salem RM, Sandholm N, Scott RA, Stolk L, Takahashi A, Tanaka T, Van’t Hooft FM, Vinkhuyzen AA, Westra HJ, Zheng W, Zondervan KT, Consortium AD, Group A-BW, Consortium CAD, Consortium CK, Glgc, Icbp, Investigators M, Mu TC, Consortium MI, Consortium P, ReproGen C, Consortium G, International Endogene C, Heath AC, Arveiler D, Bakker SJ, Beilby J, Bergman RN, Blangero J, Bovet P, Campbell H, Caulfield MJ, Cesana G, Chakravarti A, Chasman DI, Chines PS, Collins FS, Crawford DC, Cupples LA, Cusi D, Danesh J, de Faire U, den Ruijter HM, Dominiczak AF, Erbel R, Erdmann J, Eriksson JG, Farrall M, Felix SB, Ferrannini E, Ferrieres J, Ford I, Forouhi NG, Forrester T, Franco OH, Gansevoort RT, Gejman PV, Gieger C, Gottesman O, Gudnason V, Gyllensten U, Hall AS, Harris TB, Hattersley AT, Hicks AA, Hindorff LA, Hingorani AD, Hofman A, Homuth G, Hovingh GK, Humphries SE, Hunt SC, Hypponen E, Illig T, Jacobs KB, Jarvelin MR, Jockel KH, Johansen B, Jousilahti P, Jukema JW, Jula AM, Kaprio J, Kastelein JJ, Keinanen-Kiukaanniemi SM, Kiemeney LA, Knekt P, Kooner JS, Kooperberg C, Kovacs P, Kraja AT, Kumari M, Kuusisto J, Lakka TA, Langenberg C, Le Marchand L, Lehtimaki T, Lyssenko V, Mannisto S, Marette A, Matise TC, McKenzie CA, McKnight B, Moll FL, Morris AD, Morris AP, Murray JC, Nelis M, Ohlsson C, Oldehinkel AJ, Ong KK, Madden PA, Pasterkamp G, Peden JF, Peters A, Postma DS, Pramstaller PP, Price JF, Qi L, Raitakari OT, Rankinen T, Rao DC, Rice TK, Ridker PM, Rioux JD, Ritchie MD, Rudan I, Salomaa V, Samani NJ, Saramies J, Sarzynski MA, Schunkert H, Schwarz PE, Sever P, Shuldiner AR, Sinisalo J, Stolk RP, Strauch K, Tonjes A, Tregouet DA, Tremblay A, Tremoli E, Virtamo J, Vohl MC, Volker U, Waeber G, Willemsen G, Witteman JC, Zillikens MC, Adair LS, Amouyel P, Asselbergs FW, Assimes TL, Bochud M, Boehm BO, Boerwinkle E, Bornstein SR, Bottinger EP, Bouchard C, Cauchi S, Chambers JC, Chanock SJ, Cooper RS, de Bakker PI, Dedoussis G, Ferrucci L, Franks PW, Froguel P, Groop LC, Haiman CA, Hamsten A, Hui J, Hunter DJ, Hveem K, Kaplan RC, Kivimaki M, Kuh D, Laakso M, Liu Y, Martin NG, Marz W, Melbye M, Metspalu A, Moebus S, Munroe PB, Njolstad I, Oostra BA, Palmer CN, Pedersen NL, Perola M, Perusse L, Peters U, Power C, Quertermous T, Rauramaa R, Rivadeneira F, Saaristo TE, Saleheen D, Sattar N, Schadt EE, Schlessinger D, Slagboom PE, Snieder H, Spector TD, Thorsteinsdottir U, Stumvoll M, Tuomilehto J, Uitterlinden AG, Uusitupa M, van der Harst P, Walker M, Wallaschofski H, Wareham NJ, Watkins H, Weir DR, Wichmann HE, Wilson JF, Zanen P, Borecki IB, Deloukas P, Fox CS, Heid IM, O’Connell JR, Strachan DP, Stefansson K, van Duijn CM, Abecasis GR, Franke L, Frayling TM, McCarthy MI, Visscher PM, Scherag A, Willer CJ, Boehnke M, Mohlke KL, Lindgren CM, Beckmann JS, Barroso I, North KE, Ingelsson E, Hirschhorn JN, Loos RJ, Speliotes EK. Genetic studies of body mass index yield new insights for obesity biology. Nature. 2015;518:197–206.

27. Zuk O, Schaffner SF, Samocha K, Do R, Hechter E, Kathiresan S, Daly MJ, Neale BM, Sunyaev SR, Lander ES. Searching for missing heritability: Designing rare variant association studies. Proc Natl Acad Sci USA. 2014.

28. Exome Aggregation Consortium, Lek M, Karczewski K, Minikel E, Samocha K, Banks E, Fennell T, O’Donnell-Luria A, Ware J, Hill A, Cummings B, Tukiainen T, Birnbaum D, Kosmicki J, Duncan L, Estrada K, Zhao F, Zou J, Pierce-Hoffman E, Cooper D, DePristo M, Do R, Flannick J, Fromer M, Gauthier L, Goldstein J, Gupta N, Howrigan D, Kiezun A, Kurki M, Levy-Moonshine A, Natarajan P, Orozco L, Peloso G, Poplin R, Rivas M, Ruano-Rubio V, Ruderfer D, Shakir K, Stenson P, Stevens C, Thomas B, Tiao G, Tusie-Luna M, Weisburd B, Won H-H, Yu D, Altshuler D, Ardissino D, Boehnke M, Danesh J, Roberto E, Florez J, Gabriel S, Getz G, Hultman C, Kathiresan S, Laakso M, McCarroll S, McCarthy M, McGovern D, McPherson R, Neale B, Palotie A, Purcell S, Saleheen D, Scharf J, Sklar P, Sullivan PF, Tuomilehto J, Watkins H, Wilson J, Daly M, MacArthur D. Analysis of proteincoding genetic variation in 60,706 humans. Nature. 2016;536:285–291.

29. CNV Working Group of the Psychiatric Genomics Consortium, Schizophrenia Working Groups of the Psychiatric Genomics Consortium. Contribution of copy number variants to schizophrenia from a genome-wide study of 41,321 subjects. Nat Genet. 2016;49:27–35.

30. Singh T, Kurki MI, Curtis D, Purcell SM, Crooks L, McRae J, Suvisaari J, Chheda H, Blackwood D, Breen G, Pietilainen O, Gerety SS, Ayub M, Blyth M, Cole T, Collier D, Coomber EL, Craddock N, Daly MJ, Danesh J, DiForti M, Foster A, Freimer NB, Geschwind D, Johnstone M, Joss S, Kirov G, Korkko J, Kuismin O, Holmans P, Hultman CM, Iyegbe C, Lonnqvist J, Mannikko M, McCarroll SA, McGuffin P, McIntosh AM, McQuillin A, Moilanen JS, Moore C, Murray RM, Newbury-Ecob R, Ouwehand W, Paunio T, Prigmore E, Rees E, Roberts D, Sambrook J, Sklar P, St Clair D, Veijola J, Walters JT, Williams H, Swedish Schizophrenia Study, Interval Study, D. D. D. Study, UK10K Consortium, Sullivan PF, Hurles ME, O’Donovan MC, Palotie A, Owen MJ, Barrett JC. Rare loss-of-function variants in SETD1A are associated with schizophrenia and developmental disorders. Nat Neurosci. 2016;19:571–577.

31. Fromer M, Pocklington AJ, Kavanagh D, Williams H, Dwyer S, Gormley P, Georgieva L, Rees E, Palta P, Ruderfer DM, Carrera N, Humphreys I, Johnson J, Barker DD, Banks E, Milanova V, Grant SG, Hannon E, Rose SA, Chambert K, Mahajan M, Scolnick EM, Moran JL, Kirov G, Palotie A, McCarrol SA, Holmans P, Sklar P, Owen MJ, Purcell SM, O’Donovan M. De novo mutations in schizophrenia implicate synaptic networks. Nature. 2014;506:179–184.

32. Genovese G, Fromer M, Stahl EA, Ruderfer DM, Chambert K, Landen M, Moran JL, Purcell SM, Sklar P, Sullivan PF, Hultman CM, McCarrol SA. Increased burden of ultra-rare protein-altering variants among 4,877 individuals with schizophrenia. Nature Neuroscience. 2016.

33. Leonenko G, Richards AL, Walters JT, Pocklington A, Chambert K, Al Eissa MM, Sharp SI, O’Brien NL, Curtis D, Bass NJ, McQuillin A, Hultman CM, Moran JL, Sklar P, Neale BM, Holmans PA, Owen MJ, Sullivan PF, O’Donovan MC. Mutation Intolerant Genes and Targets of FMRP are Enriched for Nonsynonymous Alleles in Schizophrenia. Submitted.

34. Marouli E, Graff M, Medina-Gomez C, Lo KS, Wood AR, Kjaer TR, Fine RS, Lu Y, Schurmann C, Highland HM, Rueger S, Thorleifsson G, Justice AE, Lamparter D, Stirrups KE, Turcot V, Young KL, Winkler TW, Esko T, Karaderi T, Locke AE, Masca NG, Ng MC, Mudgal P, Rivas MA, Vedantam S, Mahajan A, Guo X, Abecasis G, Aben KK, Adair LS, Alam DS, Albrecht E, Allin KH, Allison M, Amouyel P, Appel EV, Arveiler D, Asselbergs FW, Auer PL, Balkau B, Banas B, Bang LE, Benn M, Bergmann S, Bielak LF, Bluher M, Boeing H, Boerwinkle E, Boger CA, Bonnycastle LL, Bork-Jensen J, Bots ML, Bottinger EP, Bowden DW, Brandslund I, Breen G, Brilliant MH, Broer L, Burt AA, Butterworth AS, Carey DJ, Caulfield MJ, Chambers JC, Chasman DI, Chen YI, Chowdhury R, Christensen C, Chu AY, Cocca M, Collins FS, Cook JP, Corley J, Galbany JC, Cox AJ, Cuellar-Partida G, Danesh J, Davies G, de Bakker PI, de Borst GJ, de Denus S, de Groot MC, de Mutsert R, Deary IJ, Dedoussis G, Demerath EW, den Hollander AI, Dennis JG, Di Angelantonio E, Drenos F, Du M, Dunning AM, Easton DF, Ebeling T, Edwards TL, Ellinor PT, Elliott P, Evangelou E, Farmaki AE, Faul JD, Feitosa MF, Feng S, Ferrannini E, Ferrario MM, Ferrieres J, Florez JC, Ford I, Fornage M, Franks PW, Frikke-Schmidt R, Galesloot TE, Gan W, Gandin I, Gasparini P, Giedraitis V, Giri A, Girotto G, Gordon SD, Gordon-Larsen P, Gorski M, Grarup N, Grove ML, Gudnason V, Gustafsson S, Hansen T, Harris KM, Harris TB, Hattersley AT, Hayward C, He L, Heid IM, Heikkila K, Helgeland O, Hernesniemi J, Hewitt AW, Hocking LJ, Hollensted M, Holmen OL, Hovingh GK, Howson JM, Hoyng CB, Huang PL, Hveem K, Ikram MA, Ingelsson E, Jackson AU, Jansson JH, Jarvik GP, Jensen GB, Jhun MA, Jia Y, Jiang X, Johansson S, Jorgensen ME, Jorgensen T, Jousilahti P, Jukema JW, Kahali B, Kahn RS, Kahonen M, Kamstrup PR, Kanoni S, Kaprio J, Karaleftheri M, Kardia SL, Karpe F, Kee F, Keeman R, Kiemeney LA, Kitajima H, Kluivers KB, Kocher T, Komulainen P, Kontto J, Kooner JS, Kooperberg C, Kovacs P, Kriebel J, Kuivaniemi H, Kury S, Kuusisto J, La Bianca M, Laakso M, Lakka TA, Lange EM, Lange LA, Langefeld CD, Langenberg C, Larson EB, Lee IT, Lehtimaki T, Lewis CE, Li H, Li J, Li-Gao R, Lin H, Lin LA, Lin X, Lind L, Lindstrom J, Linneberg A, Liu Y, Liu Y, Lophatananon A, Luan J, Lubitz SA, Lyytikainen LP, Mackey DA, Madden PA, Manning AK, Mannisto S, Marenne G, Marten J, Martin NG, Mazul AL, Meidtner K, Metspalu A, Mitchell P, Mohlke KL, Mook-Kanamori DO, Morgan A, Morris AD, Morris AP, Muller-Nurasyid M, Munroe PB, Nalls MA, Nauck M, Nelson CP, Neville M, Nielsen SF, Nikus K, Njolstad PR, Nordestgaard BG, Ntalla I, O’Connel JR, Oksa H, Loohuis LM, Ophoff RA, Owen KR, Packard G, Padmanabhan S, Palmer CN, Pasterkamp G, Patel AP, Pattie A, Pedersen O, Peissig PL, Peloso GM, Pennell CE, Perola M, Perry JA, Perry JR, Person TN, Pirie A, Polasek O, Posthuma D, Raitakari OT, Rasheed A, Rauramaa R, Reilly DF, Reiner AP, Renstrom F, Ridker PM, Rioux JD, Robertson N, Robino A, Rolandsson O, Rudan I, Ruth KS, Saleheen D, Salomaa V, Samani NJ, Sandow K, Sapkota Y, Sattar N, Schmidt MK, Schreiner PJ, Schulze MB, Scott RA, Segura-Lepe MP, Shah S, Sim X, Sivapalaratnam S, Small KS, Smith AV, Smith JA, Southam L, Spector TD, Speliotes EK, Starr JM, Steinthorsdottir V, Stringham HM, Stumvoll M, Surendran P, t Hart LM, Tansey KE, Tardif JC, Taylor KD, Teumer A, Thompson DJ, Thorsteinsdottir U, Thuesen BH, Tonjes A, Tromp G, Trompet S, Tsafantakis E, Tuomilehto J, Tybjaerg-Hansen A, Tyrer JP, Uher R, Uitterlinden AG, Ulivi S, van der Laan SW, Van Der Leij AR, van Duijn CM, van Schoor NM, van Setten J, Varbo A, Varga TV, Varma R, Edwards DR, Vermeulen SH, Vestergaard H, Vitart V, Vogt TF, Vozzi D, Walker M, Wang F, Wang CA, Wang S, Wang Y, Wareham NJ, Warren HR, Wessel J, Willems SM, Wilson JG, Witte DR, Woods MO, Wu Y, Yaghootkar H, Yao J, Yao P, Yerges-Armstrong LM, Young R, Zeggini E, Zhan X, Zhang W, Zhao JH, Zhao W, Zhao W, Zheng H, Zhou W, Consortium EP-I, Consortium CHDE, Exome BPC, Consortium TD-G, Go TDGC, Global Lipids Genetics C, ReproGen C, Investigators M, Rotter JI, Boehnke M, Kathiresan S, McCarthy MI, Willer CJ, Stefansson K, Borecki IB, Liu DJ, North KE, Heard-Costa NL, Pers TH, Lindgren CM, Oxvig C, Kutalik Z, Rivadeneira F, Loos RJ, Frayling TM, Hirschhorn JN, Deloukas P, Lettre G. Rare and low-frequency coding variants alter human adult height. Nature. 2017;542:186–190.

35. Purcell SM, Moran JL, Fromer M, Ruderfer D, Solovieff N, Roussos P, O’Dushlaine C, Chambert K, Bergen K, Kahler A, Duncan LH, Stahl E, Genovese G, Fernandez E, Collins MO, Komiyama NH, Choudhary JS, Magnusson PKE, Banks E, Shakir K, Garimella K, Fennell T, de Pristo M, Grant SGN, Haggarty SJ, Garbriel S, Scolnick EM, Lander ES, Hultman CM, Sullivan PF, McCarrol SA, Sklar P. A polygenic burden of rare disruptive mutations in schizophrenia. Nature. 2014;506:185–190.

36. Cross-Disorder Group of the Psychiatric Genomics Consortium. Identification of risk loci with shared effects on five major psychiatric disorders: a genome-wide analysis. Lancet. 2013;381:1371–1379.

37. Ripke S, O’Dushlaine C, Chambert K, Moran JL, Kähler A, Akterin S, Bergen S, Collins AL, Crowley J, Fromer M, Kim Y, Lee SH, Magnusson PK, Sanchez N, Stahl EA, Williams S, Wray N, Xia K, Bettella F, Børglum AD, Bulik-Sullivan BK, Cormican P, Craddock N, de Leeuw C, Durmishi N, Gill M, Golimbet V, Hamshere ML, Holmans P, Hougaard DM, Kendler KS, Lin K, Morris DW, Mors O, Mortensen PB, Neale B, O’Neill FA, Owen MJ, Pejovic Milovancevic M, Posthuma D, Powell J, Richards AL, Riley BP, Ruderfer D, Rujescu D, Sigurdsson E, Silagadze T, Smit AB, Stefansson H, Steinberg S, Suvisaari J, Tosato S, Walters JT, Verhage M, Multicenter Genetic Studies of Schizophrenia Consortium, Psychosis Endophenotypes Consortium, Wellcome Trust Case-Control Consortium2, Bramon E, Corvin AP, O’Donovan MC, Stefansson K, Scolnick E, Purcell S, McCarroll S, Sklar P, Hultman C, Sullivan PF. Genome-wide association analysis identifies 13 new risk loci for schizophrenia. Nature Genetics. 2013;45:1150–1159.

38. Breen G, Li Q, Roth BL, O’Donnell P, Didriksen M, Dolmetsch R, O’Reilly PF, Gaspar HA, Manji H, Huebel C, Kelsoe JR, Malhotra D, Bertolino A, Posthuma D, Sklar P, Kapur S, Sullivan PF, Collier DA, Edenberg HJ. Translating genome-wide association findings into new therapeutics for psychiatry. Nat Neurosci. 2016;19:1392–1396.

39. Gaspar HA, Breen G. Pathways analyses of schizophrenia GWAS focusing on known and novel drug targets. Submitted.

40. Nelson MR, Tipney H, Painter JL, Shen J, Nicoletti P, Shen Y, Floratos A, Sham PC, Li MJ, Wang J, Cardon LR, Whittaker JC, Sanseau P. The support of human genetic evidence for approved drug indications. Nat Genet. 2015;47:856–860.

41. Franke B, Stein J, Ripke S, Anttila V, Hibar DP, van Hulzen KJ, Vasquez AA, Smoller JW, Nichols TE, Neale MC, McIntosh A, Lee PC, McMahon F, Meyer-Lindenberg A, Mattheisen M, Andreassen O, Gruber O, Sachdev PS, Roiz-Santianez R, Saykin AJ, Ehrlich S, Mather KA, Turner J, Schwartz E, Thalamuthu A, Yoo Y, Martin NG, Wright MJ, Schizophrenia Working Group of the Psychiatric Genomics Consortium, ENIGMA Consortium, O’Donovan M C, Thompson P, Neale B, Medland SE, Sullivan PF. Genetic influences on schizophrenia and subcortical brain volumes: large-scale proof-of-concept and roadmap for future studies. Nature Neuroscience. 2016;19:420–431.

42. Domingue BW, Belsky D, Conley D, Harris KM, Boardman JD. Polygenic Influence on Educational Attainment: New evidence from The National Longitudinal Study of Adolescent to Adult Health. AERA Open. 2015;1:1–13.

43. Zheng J, Erzurumluoglu AM, Elsworth BL, Kemp JP, Howe L, Haycock PC, Hemani G, Tansey K, Laurin C, Early G, Lifecourse Epidemiology Eczema C, Pourcain BS, Warrington NM, Finucane HK, Price AL, Bulik-Sullivan BK, Anttila V, Paternoster L, Gaunt TR, Evans DM, Neale BM. LD Hub: a centralized database and web interface to perform LD score regression that maximizes the potential of summary level GWAS data for SNP heritability and genetic correlation analysis. Bioinformatics. 2017;33:272–279.

44. Roussos P, Mitchell AC, Voloudakis G, Fullard JF, Pothula VM, Tsang J, Stahl EA, Georgakopoulos A, Ruderfer DM, Charney A, Okada Y, Siminovitch KA, Worthington J, Padyukov L, Klareskog L, Gregersen PK, Plenge RM, Raychaudhuri S, Fromer M, Purcell SM, Brennand KJ, Robakis NK, Schadt EE, Akbarian S, Sklar P. A role for noncoding variation in schizophrenia. Cell reports. 2014;9:1417–1429.

45. Won H, de la Torre-Ubieta L, Stein JL, Parikshak NN, Huang J, Opland CK, Gandal MJ, Sutton GJ, Hormozdiari F, Lu D, Lee C, Eskin E, Voineagu I, Ernst J, Geschwind DH. Chromosome conformation elucidates regulatory relationships in developing human brain. Nature. 2016;538:523–527.

46. Pathway Analysis Subgroup of the Psychiatric Genomics Consortium. Psychiatric genome-wide association study analyses implicate neuronal, immune and histone pathways. Nat Neurosci. 2015;18:199–209.

47. Finucane HK, Bulik-Sullivan BK, Gusev A, Trynka G, Reshef Y, Loh P-R, Anttilla V, Xu H, Zang C, Farh K, Ripke S, Day F, ReproGen Consortium, Schizophrenia Working Group of the Psychiatric Genomics Consortium, RACI Consortium, Purcell S, Stahl EA, Lindstrom S, Perry JRB, Okada Y, Raychaudhuri S, Daly M, Patterson N, Neale BM, Price AL. Partitioning heritability by functional category using GW AS summary statistics. Nature Genetics. 2015;47:1228–1235.

48. Fromer M, Roussos P, Sieberts SK, Johnson JS, Kavanagh DH, Perumal T, Ruderfer DM, Shah HR, Klei LL, Kramer R, Pinto D, Gümüs ZH, Clcek AE, Dang KK, Browne A, Lu C, Readhead B, Wang Y-C, Mahajan MC, Derry JMJ, Dudley J, Logsdon BA, Talbot K, Zhu J, Zhang B, Sullivan PF, Chess A, Purcell SM, CommonMind Consortium, Shinobu LA, Mangravite LM, Toyoshiba H, Gur RE, Hahn C-G, Lewis DA, Haroutunian V, Peters MA, Lipska BK, Buxbaum JD, Schadt EE, Hirai K, Domenici E, Devlin B, Sklar P. Gene expression elucidates functional impact of polygenic risk for schizophrenia. Nature Neuroscience. 2016;19:1442–1453.

49. Akbarian S, Liu C, Knowles JA, Vaccarino FM, Farnham PJ, Crawford GE, Jaffe AE, Pinto D, Dracheva S, Geschwind DH, Mill J, Nairn AC, Abyzov A, Pochareddy S, Prabhakar S, Weissman S, Sullivan PF, State MW, Weng Z, Peters MA, White KP, Gerstein MB, Amiri A, Armoskus C, Ashley-Koch AE, Bae T, Beckel-Mitchener A, Berman BP, Coetzee GA, Coppola G, Francoeur N, Fromer M, Gao R, Grennan K, Herstein J, Kavanagh DH, Ivanov NA, Jiang Y, Kitchen RR, Kozlenkov A, Kundakovic M, Li M, Li Z, Liu S, Mangravite LM, Mattei E, Markenscoff-Papadimitriou E, Navarro FC, North N, Omberg L, Panchision D, Parikshak N, Poschmann J, Price AJ, Purcaro M, Reddy TE, Roussos P, Schreiner S, Scuderi S, Sebra R, Shibata M, Shieh AW, Skarica M, Sun W, Swarup V, Thomas A, Tsuji J, van Bakel H, Wang D, Wang Y, Wang K, Werling DM, Willsey AJ, Witt H, Won H, Wong CC, Wray GA, Wu EY, Xu X, Yao L, Senthil G, Lehner T, Sklar P, Sestan N. The PsychENCODE project. Nat Neurosci. 2015;18:1707–1712.

50. Gilissen C, Hehir-Kwa JY, Thung DT, van de Vorst M, van Bon BW, Willemsen MH, Kwint M, Janssen IM, Hoischen A, Schenck A, Leach R, Klein R, Tearle R, Bo T, Pfundt R, Yntema HG, de Vries BB, Kleefstra T, Brunner HG, Vissers LE, Veltman JA. Genome sequencing identifies major causes of severe intellectual disability. Nature. 2014;511:344–347.

51. Sanders SJ, He X, Willsey AJ, Ercan-Sencicek AG, Samocha KE, Cicek AE, Murtha MT, Bal VH, Bishop SL, Dong S, Goldberg AP, Jinlu C, Keaney JF, 3rd, Klei L, Mandell JD, Moreno-De-Luca D, Poultney CS, Robinson EB, Smith L, Solli-Nowlan T, Su MY, Teran NA, Walker MF, Werling DM, Beaudet AL, Cantor RM, Fombonne E, Geschwind DH, Grice DE, Lord C, Lowe JK, Mane SM, Martin DM, Morrow EM, Talkowski ME, Sutcliffe JS, Walsh CA, Yu TW, Autism Sequencing C, Ledbetter DH, Martin CL, Cook EH, Buxbaum JD, Daly MJ, Devlin B, Roeder K, State MW. Insights into Autism Spectrum Disorder Genomic Architecture and Biology from 71 Risk Loci. Neuron. 2015;87:1215–1233.

52. Devlin B, Scherer SW. Genetic architecture in autism spectrum disorder. Curr Opin Genet Dev. 2012;22:229–237.

53. Levinson DF, Duan J, Oh S, Wang K, Sanders AR, Shi J, Zhang N, Mowry BJ, Olincy A, Amin F, Cloninger CR, Silverman JM, Buccola NG, Byerley WF, Black DW, Kendler KS, Freedman R, Dudbridge F, Pe’er I, Hakonarson H, Bergen SE, Fanous AH, Holmans PA, Gejman PVCopy number variants in schizophrenia: Confirmation of five previous findings and new evidence for 3q29 microdeletions and VIPR2 duplications. Am J Psychiatry. 2011;168:302–316.

